# PglZ from Type I BREX phage defence systems is a metal-dependent nuclease that forms a sub-complex with BrxB

**DOI:** 10.1101/2025.03.26.645558

**Authors:** Jennifer J. Readshaw, Lindsey A. Doyle, Maria Puiu, Abigail Kelly, Andrew Nelson, Alex J. Kaiser, Sydney McGuire, Julieta Peralta-Acosta, Darren L. Smith, Barry L. Stoddard, Brett K. Kaiser, Tim R. Blower

**Affiliations:** Department of Biosciences, Durham University, Stockton Road, Durham, DH1 3LE, UK; Division of Basic Sciences, Fred Hutchinson Cancer Center, 1100 Fairview Ave. N. Seattle WA 98019, USA; Department of Applied Sciences, University of Northumbria, Newcastle Upon Tyne NE1 8ST, UK; Department of Biology, Seattle University, 901 12th Ave. Seattle WA 98122, USA; New England Biolabs, 240 County Road, Ipswich, MA 01938, USA

**Author notes:** These authors contributed equally (Jennifer J. Readshaw, Lindsey A. Doyle).

**Keywords:** Phage defence, Bacteriophage exclusion, nuclease, metal-dependent, BREX

## Abstract

BREX (*B*acte*r*iophage *Ex*clusion) systems, identified through shared identity with Pgl (*P*hage *G*rowth *L*imitation) systems, are a widespread, highly diverse group of phage defence systems found throughout bacteria and archaea. The varied BREX Types harbour multiple protein subunits (between four and eight) and all encode a conserved putative phosphatase (PglZ aka BrxZ) and an equally conserved, putative ATPase (BrxC). Almost all BREX systems also contain a site-specific methyltransferase (PglX aka BrxX). Despite having determined the structure and fundamental biophysical and biochemical behaviours for the PglX methyltransferase, the BrxL effector, the BrxA DNA-binding protein and the BrxR transcriptional regulator, the mechanism by which BREX impedes phage replication remains largely undetermined. In this study, we identify a stable BREX sub-complex of PglZ:BrxB, validate the structure and dynamic behaviour of that sub-complex, and assess the biochemical activity of PglZ, revealing it to be a metal-dependent nuclease. PglZ can cleave cyclic oligonucleotides, linear oligonucleotides, plasmid DNA and both non-modified and modified linear phage genomes. PglZ nuclease activity has no obvious role in BREX-dependent methylation, but does contribute to BREX phage defence. BrxB binding does not impact PglZ nuclease activity. These data contribute to our growing understanding of the BREX phage defence mechanism.

## Introduction

Up to 10% of bacterial and archaeal genes are dedicated to phage defence (1). The mechanisms employed to defend against phage infection are diverse and include prevention of viral entry, induction of cell dormancy or death upon infection, and mechanisms that degrade viral genomes or block viral DNA replication (2, 3). Phages combat these systems through the evolution of elaborate countermeasures that block their action, leading to viral resistance and a continuous arms race between phage and bacterial populations (4). Recent analyses have demonstrated that bacteria encode far more phage defence systems than just the most well-studied ‘first responder’ systems such as restriction endonucleases and CRISPR (5–10). Furthermore, many of these newly discovered bacterial systems display obvious similarities to human innate viral defense systems, implying common evolutionary origins and related mechanisms of action (11).

Originally discovered in the early 1980s (12), *P*hage *G*rowth *L*imitation (Pgl) (13, 14) and related *B*acte*r*iophage *Ex*clusion (BREX) systems are widespread in bacterial and archaeal species (15). BREX systems are encoded by single operons, often within genetic defence islands, and are currently categorised into at least six types based on the number of genes in each system (typically four to eight) and on the sequence-based functional annotation of those individual genes and putative translated protein subunits (15); Type I systems, the most common subtype, comprise six conserved genes and can readily be assayed for the two phenotypes of phage defence and BREX-dependent methylation though the mechanisms are unknown.

Pgl and BREX systems have two genes in common. The first is named *pglZ* (aka *brxZ*) and the second is *brxC* (15). The PglZ domain of a two-component signalling system response regulator, PorX, has recently been shown to degrade cyclic nucleotides (16), but any equivalent activity within BREX has not yet been explored. Beyond PglZ and BrxC, most BREX systems include a gene encoding a site-specific methyltransferase, termed PglX (aka BrxX). The identity and order of the remaining genes in each BREX type vary significantly: various BREX systems encode protein subunits with domains that display recognisable homology to kinases, phosphatases, DNA and/or nucleotide binding domains, DNA modification enzymes, chambered AAA+ ATPases, and/or DNA helicases. Several BREX subunits are quite large, with significant regions of unknown structure-function properties and behaviours flanking domains with well-annotated putative functions. Type I BREX systems, like their related counterparts, do not contain any readily identifiable DNA nuclease domains or subunits and appear to likely restrict phage by inhibiting phage DNA replication within the infected bacterial cell.

Despite having previously determined the high resolution structures and biochemical activities of the PglX methyltransferase (17, 18), BrxR (a WYL-domain helix-turn-helix DNA binding transcriptional regulator) (19, 20), BrxA (a small DNA binding protein) (21), and BrxL (a chambered AAA^+^ ATPase and dsDNA binding protein) (22), the mechanism by which BREX systems function to restrict phage replication and to protect the host genome from the system’s activity is still largely unknown. We have performed analyses to further characterise several Type I BREX systems we have previously investigated: those from *Salmonella* Typhimurium (17, 23) and *Escherichia fergusonii* (24) (**Fig. 1A**), and from *Acinetobacter* (20, 22). Using these systems, *in vivo* pull-down and co-expression analysis identified larger BREX complexes and a stable sub-complex formed by PglZ and BrxB. Computational models of the PglZ:BrxB interactions and their likely conformation and dynamic behaviour when bound to one another was validated through single-particle cryoEM analyses. Subsequent biochemical analysis has identified that PglZ recapitulates PorX activity and is a metal-dependent nuclease that can cleave not only a broad range of cyclic and linear oligonucleotides, but also plasmid and linear dsDNA. The BrxB interaction does not impact PglZ nuclease activity, nor is a nuclease required for BREX-dependent methylation. Nuclease activity does, however, contribute to BREX phage defence. These data contribute to our growing understanding of the elusive BREX mechanism.

**Figure 1.**
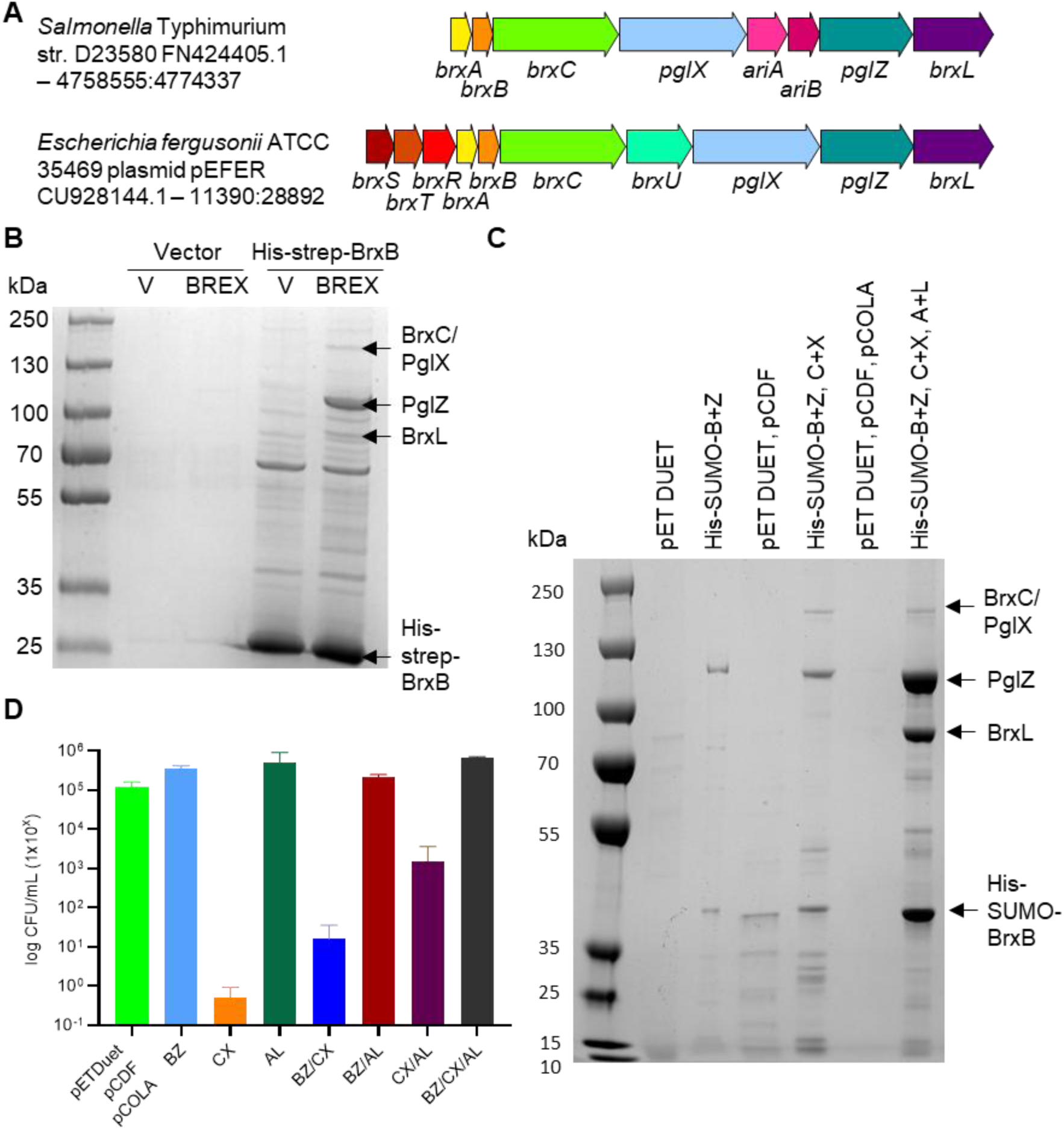
BREX proteins form higher order complexes. (**A**) Schematic of the BREX phage defence islands from *Salmonella* and *Escherichia fergusonii*. The *Salmonella* island encodes BREX and PARIS (*ariA*, *ariB*) defence systems. The *E. fergusonii* island encodes BREX and a Type IV restriction enzyme of the GmrSD family, BrxU. (**B**) Pull-down of *Salmonella* BREX complexes. *E. coli* BL21 (DE3) pRARE was transformed with an inducible plasmid expressing His-strep tagged BrxB (or vector control), and a second plasmid expressing the six BREX genes *brxA*, *brxB*, *brxC*, *pglX*, *pglZ* and *brxL* (or vector control). Pull-down samples were analysed by SDS-PAGE and indicated bands were identified by mass spectrometry. (**C**) Expression of *Salmonella* BREX proteins in pairs from pET DUET-based vectors, pulled down with His-SUMO-BrxB. Pull-down samples were analysed on SDS-PAGE and indicated bands were identified by mass spectrometry. (**D**) Toxicity during expression of *Salmonella* BREX proteins, measured as viable counts. Error bars represent the standard deviation of the mean from triplicate data.

## Materials and Methods

### Bacterial strains and culture conditions

*E. coli* strains DH5α (Invitrogen), ER2796 (New England Biolabs) (25), ER2566 (New England Biolabs), Rosetta 2 (DE3) pLysS (Novagen), BL21 (DE3) pRARE (Novagen), BL21 (DE3) RIL (Novagen) and T7 Express (New England Biolabs) were routinely grown at 37 °C, either on agar plates or shaking at 150 rpm for liquid cultures. 2x Yeast Extract Tryptone (YT) was used as the standard growth media for liquid cultures, and Luria Broth (LB) was supplemented with 0.35% (w/v) or 1.5% (w/v) agar for semi-solid and solid agar plates, respectively. When necessary, growth media was supplemented with ampicillin (Ap, 100 μg/ml), chloramphenicol (Cm, 25 μg/ml), kanamycin (Km, 100 μg/mL, spectinomycin (Sp, 100 μg/mL), isopropyl-β-D-thiogalactopyranoside (IPTG, 1 mM). Growth was monitored using a spectrophotometer (WPA Biowave C08000) measuring optical density at 600 nm (OD_600_).

### DNA isolation and manipulation

Plasmid DNA was purified from transformed DH5α cells using an NEB Monarch® Plasmid MiniPrep kit following the manufacturer’s instructions. Larger amounts of negatively supercoiled plasmid pSG483 (26) DNA for assays was purified from transformed DH5α cells using a Machery-Nagel NucleoBond Xtra Midi Plus EF kit following the manufacturer’s instructions. Plasmid DNA was eluted in MiliQ and stored at −20 °C. Plasmids are described in **Supplementary Table S1**.

Phage genomic DNA was purified by incubating 450 μl phage lysate with 4.5 μl DNase I (1 mg/ml; Sigma-Aldrich) and 12.5 μl RNase A (10 mg/ml; ThermoFisher) for 30 min at 37 °C. The lysate was further incubated with 2.25 μl proteinase K (20 mg/ml; Sigma-Aldrich) and 23 μl of 10% (w/v) SDS for 30 min at 37 °C. The sample was mixed with 500 μl UltraPure™ phenol:chloroform:isoamyl alcohol (25:24:1; v/v/v) (ThermoFisher) and centrifuged at 16,000 x *g* for 5 min at 4 °C. The aqueous layer was removed carried forward, and the previous step was repeated. The resulting aqueous layer was mixed with 500 μl chloroform:isoamyl alcohol (24:1; v/v) and centrifuged at 16,000 x *g* for 5 min at 4 °C. The aqueous layer was carried forward and incubated with 45 μl 3 M sodium acetate pH 5.2 and 500 μl isopropanol for 15 min at room temperature, before being centrifuged at 16,000 x *g* for 20 min and 4 °C. The supernatant was removed, and the pellet was washed with 70% ethanol by gentle aspiration before being dried at room temperature. The dry pellet was soaked in 50 μl of MiliQ and incubated overnight at 4 °C. The gDNA was analysed on a 0.75% 1x TAE agarose gel by agarose gel electrophoresis, and stored at −20 °C.

### Preparation of nicked, linear, and relaxed form pSG483

Linear pSG483 was obtained through incubation of 10 μg pSG483 with 10 units of BamHI-HF® (New England Biolabs) in 1x CutSmart buffer (New England Biolabs) for 1 h at 37 °C. The enzyme was deactivated by incubation at 65 °C for 10 min. Nicked pSG483 was obtained by incubating 10-50 μg pSG483 with 10 units Nb.Bpu10I (ThermoFisher) in 1x Buffer R (ThermoFisher) for 4 h at 37 °C. The reaction was terminated by incubation at 80 °C for 20 min.

For production of relaxed pSG483, 50 μg nicked pSG483 was further incubated with 1 mM ATP and 10 units T4 ligase (New England Biolabs) for 16 h at room temperature. After ligation, an equal volume of UltraPure™ phenol:chloroform:isoamyl alcohol (25:24:1; v/v/v) (ThermoFisher) was added to the reaction mixture, vortexed briefly, and centrifuged at 16,000 x *g* for 2 min. The resulting aqueous layer was removed and carried forward. An equal volume of chloroform was added to the aqueous layer before centrifugation at 16,000 x *g* for 2 min. The resulting aqueous layer was carried forward and 1/10 volume of 3 M sodium acetate pH 5.2 was added, followed by 2 volumes of 100% ethanol. The sample was mixed by pipetting and stored at −80 °C for 30 min. The sample was centrifuged at 16,000 x *g* for 20 min at 4 °C. The ethanol was removed, and the DNA pellet dried at room temperature. The DNA pellet was resuspended to 300 ng/μl with MiliQ. All DNA products were analysed by agarose gel electrophoresis prior to storage at −20 °C.

### Bacterial growth assays

T7 Express *E. coli* cells were transformed with: empty vectors pETDuet, pCDFDuet and pCOLADuet (i), pTRB710 (His-SUMO-BrxB and PglZ (BZ)) (ii), pTRB759 (BrxC and PglX (CX)) (iii), pTRB758 (BrxA and BrxL (AL)) (iv) and combinations of BZ/CX (v), BZ/AL (vi), CX/AL (vii) and BZ/CX/AL (viii). Colonies were inoculated and grown overnight in 5 ml 2x YT with respective antibiotics at 37 °C shaking at 180 rpm. Cultures were re-seeded 1:100 (v/v) in 100 ml 2x YT with the relevant antibiotics and grown at 37 °C until OD_600_ reached ∼0.4. ODs of all cultures were then normalised to OD_600_ of ∼1.0. Cultures were then serially diluted 10^−1^ to 10^−7^ and spotted on LB agar plates containing the relevant antibiotics +/− IPTG for induction of each complex combination. Plates were then incubated overnight at 37 °C and imaged for colony counting and CFU/ml determination.

### Protein expression and purification

For large-scale expression of *Salmonella* PglZ or co-expression of *Salmonella* His-SUMO-BrxB and PglZ, *E. coli* ER2566 was transformed with pSALMZ and pTRB710, respectively. For large-scale expression of *E. fergusonii* proteins for biochemistry, *E. coli* ER2566 was transformed with pTRB449 (PglZ) and pTRB444 (BrxB). *E. coli* Rosetta (DE3) pLysS was also transformed with pTRB444. *E. fergusonii* mutant derivatives were expressed by transforming *E. coli* ER2566 with plasmids pTRB729 (PglZ T538A), pTRB730 (PglZ H741A), pTRB763 (PglZ T538A/H741A), pTRB727 (BrxB W135A), and pTRB726 (BrxB R46A), and transforming *E. coli* Rosetta (DE3) pLysS with pTRB724 (BrxB E47A), pTRB725 (BrxB S133A), pTRB726 (BrxB R46A), pTRB727 (BrxB W135A), and pTRB728 (BrxB E89A).

The same procedures were used for both *Salmonella* and *E. fergusonii* proteins. Single colonies were used to inoculate 70 ml 2x YT for overnight growth at 37 °C shaking at 180 rpm. Starter cultures were re-seeded 1:100 (v/v) into 1 L 2x YT containing the relevant antibiotic(s) in 2 L baffled flasks and incubated at 37 °C until the OD_600_ reached ∼0.4. At this point, the incubation temperature was reduced to 18 °C for overnight incubation and expression was induced with IPTG. Cells were harvested by centrifugation at 4,200 x *g* for 20 min at 4 °C. Cell pellets were resuspended on ice in ice-cold A500 (20 mM Tris HCl pH 7.9, 500 mM NaCl, 10% (v/v) glycerol, 10 mM imidazole). Resuspended cells were disrupted by sonication (45% amplitude, 10 s on 20 s off pulse intervals, 2 min) and clarified by centrifugation at 45,000 x *g* for 45 min at 4 °C. Clarified cell lysate was loaded onto a 5 ml HisTrap HP column (Cytiva) pre-equilibrated in A100 (20 mM Tris HCl pH 7.9, 100 mM NaCl, 10% (v/v) glycerol, 10 mM imidazole). The HisTrap column was then washed with 50 ml A100, and bound proteins were eluted directly onto a pre-equilibrated 5 ml HiTrap Q HP column using B100 (20 mM Tris HCl pH 7.9, 100 mM NaCl, 10% (v/v) glycerol, 250 mM imidazole). The Q HP column was washed with 50 ml A100 and transferred to an ÅktaÔ Pure (Cytiva), and the target protein was eluted by anion exchange chromatography (AEC) using a salt gradient from 100% A100 to 60% C1000 (20 mM Tris HCl pH 7.9, 1 M NaCl, 10% (v/v) glycerol). Chromatographic peak fractions were collected, pooled, and incubated overnight in the presence of human sentrin/SUMO-specific protease 2 (hSENP2) to facilitate the cleavage of the His-SUMO tag at 4 °C. The following day, the SENP-treated sample was applied to a second His-Trap HP column pre-equilibrated in A100. The flow-through containing untagged target protein was collected and concentrated by centrifugation using the appropriate MWCO Vivaspin concentrator (Sartorius). Concentrated protein samples were applied to a HiPrep™ 16/60 Sephacryl® S-200 HR column (S-200; Cytiva) pre-equilibrated with 1.2 column volumes (CV) of sizing buffer (50 mM Tris HCl pH 7.9, 500 mM KCl, 10% (v/v) glycerol) for further purification by size exclusion chromatography (SEC). SEC peak fractions were pooled and analysed by SDS-PAGE, then concentrated as described previously and quantified using a NanoDrop 2000 Spectrophotometer (Thermo Fisher). Final purified samples for biochemical analysis were resuspended in a 1:2 mixture of protein sample:storage buffer (50 mM Tris HCl pH 7.9, 500 mM KCl, 70% (v/v) glycerol) and flash frozen in liquid nitrogen for storage at −80 °C.

*Acinetobacter* proteins were expressed as follows: plasmids encoding PglZ^Aci^ with a C-terminal, thrombin-cleavable Twin-Strep Tag (pET15b.PglZ-TST) and untagged BrxB^Aci^ (pET24d.BrxB) were used to co-transform *E. coli* BL21 (DE3) RIL. Overnight cultures (25 ml, LB) were diluted 100-fold into 1 L of LB and grown to an OD_600_ of 0.6, at which time IPTG was added to 200 µM. Cultures were incubated at 16 °C for 18 h, pelleted by centrifugation at 4,200 x *g* for 45 min, and the pellets were stored at −20 °C. Pellets were lysed in Buffer W (100 mM Tris-HCl pH 8.0, 150 mM NaCl, 1 mM EDTA), centrifuged for 25 min at 18,000 x *g* in an SS34 rotor at 4 °C, and the supernatant was filtered through a 5 µm syringe filter. The soluble lysate was bound to streptactin resin, washed with 10 CV of Buffer W and eluted in Buffer E (100 mM Tris-HCl pH 8.0, 150 mM NaCl, 1 mM EDTA, 2.5 mM desthiobiotin). Samples were concentrated in a 30 kDa MWCO Amicon filter (EMD Millipore) and purified by SEC on a SEC650 column (BioRad) equilibrated in 25 mM Tris-HCl pH 7.5, 200 mM NaCl.

### SDS-PAGE electrophoresis

Protein samples were analysed by SDS-PAGE. For protein purity analysis, 4 μg of PglZ and derivative mutants, and 4 μg of BrxB and derivative mutants were made up to 10 μl with A100 and mixed with 5 μl 3x sample buffer (187.5 mM Tris HCl pH 6.8, 6% (w/v) SDS, 30% (v/v) glycerol, 0.03% (w/v) bromophenol blue, 150 mM DTT) and denatured for 10 min at 95 °C. Protein samples were loaded onto and resolved in 15% (v/v) and 12% (v/v) poly-acrylamide gels, respectively, in 1x Tris-glycine running buffer (25 mM Tris, 192 mM glycine, 0.1% (w/v) SDS) at 180 V for 1h 15 min. For PglZ and BrxB interaction analysis post analytical SEC, 30 μl of fractions for analysis were mixed with 6 μl of 6x sample buffer (375 mM Tris HCl pH 6.8, 12% (w/v) SDS, 60% (v/v) glycerol, 0.06% (w/v) bromophenol blue, 300 mM DTT) and resolved in 15% (v/v) poly-acrylamide gels as described previously. Gels were stained with Quick Coomassie (Protein Ark) and destained with MiliQ. Gel images were obtained on a ChemiDoc™ Imaging System on the Coomassie brilliant blue setting (BioRad).

### Protein pull-down assays

His-strep tagged BrxB (expressed from 2HR-T, addgene #29718) was used as bait for pull-down assays of BREX components expressed from pCOLA DUET1 in *E. coli* BL21 (DE3) pRARE. Overnight cultures were used to inoculate 25 ml of 2xYT with the relevant antibiotics to OD 0.1 before growth at 37 °C 180 rpm to OD ∼0.8. Cultures were induced with 1 mM IPTG and incubated at 18 °C with shaking overnight. Cells were harvested at 4,200 x *g* for 15 min before freezing at −70 °C. Pellets were defrosted and resuspended in 10 ml 100 mM Tris pH 7.9 150 mM NaCl before sonication (5 min of 10 s pulses at 30% power). Lysates were clarified by centrifugation at 45,000 x *g* for 10 min at 4 °C. Clarified lysates were incubated with 200 µl pre-equilibrated Strep-Tactin Sepharose High Performance resin (Cytiva) at 4 °C for 90 min with rolling before application to a Proteus Mini Spin column (ProteinArk). The resin was washed three times with 100 mM Tris pH 7.9 150 mM NaCl, before incubation of the resin in the column with 50 µl 100 mM Tris pH 7.9 150 mM NaCl, 2.5 mM desthiobiotin. The protein was eluted from the column by centrifugation at 12,000 x *g* for 1 min. The 50 µl eluate was re-applied and re-incubated with the resin before a second centrifugal elution step. Pull-down products were separated and visualised on a 4-15% SDS-PAGE gel.

His-SUMO (27) tagged BrxB was used as bait for pull-down assays following induced expression of all BREX components. Plasmids were co-expressed in T7 *E. coli* cells for protein complex formation in the following way: empty vector pETDuet1 as control for His-SUMO-BZ (i), empty vectors pETDuet1 and pCDFDuet1 as control for BZ/CX (ii), empty vectors pETDuet1, pCDFDuet1 and pCOLADuet1 as control for BZ/CX/AL (iii) BZ on its own (iv), BZ along with CX (v) and BZ with CX and with AL (vi). Single colonies were inoculated in 20 ml 2x YT for overnight growth at 37 °C shaking at 180 rpm. Started cultures were then re-seeded into 1 L 2x YT containing the relevant antibiotic(s) in 2 L baffled flasks and incubated at 37 °C until the OD_600_ reached ∼0.4. Cultures were then incubated at 18 °C overnight and expression was induced with IPTG. Cells were harvested by centrifugation at 4,200 x *g* for 20 min at 4 °C. Cell pellets were resuspended on ice in ice-cold A500. Resuspended cells were disrupted by sonication (45% amplitude, 5 s on 10 s off pulse intervals, 5 min) and centrifuged at 45,000 x *g* for 30 min at 4 °C. Clarified cell lysate was loaded onto a 5 ml HisTrap HP column (Cytiva) pre-equilibrated in A100. The HisTrap column was then washed with 50 ml A100, and transferred to an ÅktaÔ Pure (Cytiva) for complex elution using an imidazole gradient from 10 mM imidazole to 250 mM using B100. Peak fractions were collected accordingly and concentrated by centrifugation using the MWCO Vivaspin concentrator (Sartorius) of appropriate size. Concentrated complexes were then loaded on a HiPrep™ 16/60 Sephacryl® S-200 HR column (S-200; Cytiva) pre-equilibrated with 1.2 CV of sizing buffer for size exclusion chromatography (SEC) purification. SEC peak fractions were pooled and analysed by SDS-PAGE, then concentrated and quantified using a NanoDrop 2000 Spectrophotometer (Thermo Fisher).

### Bis(4-nitrophenyl) phosphate phosphodiesterase activity assay

EDTA treated PglZ (PglZ EDTA) was prepared by incubating PglZ with 1 mM EDTA for 15 min at room temperature. The EDTA was removed by centrifuging in a 30 kDa MWCO Vivaspin™ ultrafiltration spin column (Cytiva) at 12,000 x *g* at 4 °C, until the volume < 100 μl. The sample was resuspended in ∼400 μl A100 and centrifuged again. This was repeated twice. PglZ and PglZ EDTA (2.2 μM) were incubated with 1x PglZ buffer (“ZB”: 50 mM Tris HCl pH 8.0, 150 mM NaCl) with (PglZ EDTA) or without 0.5 mM MgCl_2_, MnCl_2_, or CaCl_2_, for 30 mins at room temperature. The phosphodiesterase reaction was initiated by adding 10 μl 25 mM bis(4-nitrophenyl)phosphate (bis-*p*NPP, Merck) to 90 μl PglZ reaction mix, and monitoring the release of reaction product, *p*-nitrophenol, for 2 hours at 37 °C by measuring the absorbance at 405 nm on a SPECTROstar® *Nano* microplate reader (BMG Labtech). PglZ derived mutants were also assayed for activity as described. Triplicate reactions were performed per assay, and the assay was completed in triplicate. Control reactions comprised 1x ZB with and without 0.5 mM MgCl_2_, MnCl_2_, or CaCl_2_ in the presence of bis-*p*NPP.

### Nucleotide cleavage assay

PglZ and derivative mutants (2 μM) were incubated with 10 μM ZnCl_2_ or MnCl_2_ in 1x ZB with 10 μM of the following nucleotides: cyclic hexa-adenosine monophosphate (cA6); cyclic tetra-adenosine monophosphate (cA4); cyclic tri-adenosine monophosphate (cA3); 3′,5′-cyclic di-adenylate (cA2); 5′-phosphoadenylyl-(3′-5′)-adenosine (pApA); 5′-phosphoadenylyl-(3′-5′)-guanosine (pApG); 5′-phosphoguanylyl-(3′-5′)-guanosine (pGpG); 3′,5′-cyclic di-guanylate (cG2); 3′,5′-cyclic adenosine monophosphate (cAMP); 3′,5′-cyclic uridine monophosphate (cUMP); 3′,5′-cyclic thymidine monophosphate (cTMP); 3′,5′-cyclic cytidine monophosphate (cCMP); 3′,5′-cyclic guanosine monophosphate (cGMP); 2′3′-cyclic uridine monophosphate (2′3′ cUMP); 2′3′-cyclic adenosine monophosphate (2′3′ cAMP); 2′3′-cyclic guanosine monophosphate (2′3′ cGMP); cyclic adenosine-(3′-5′)-monophosphate adenosine-(3′-5′)-monophosphate guanosine-(3′-5′)-monophosphate (c[A(3′5′)pA(3′5′)pG(3′5′)p]); cyclic adenosine-(2′-5′)-monophosphate guanosine-(3′-5′)-monophosphate (c[A(2′5′)pG(3′5′)p]); cyclic adenosine-(3′-5′)-monophosphate guanosine-(3′-5′)-monophosphate (c-ApGp); P^1^-(5′-adenosyl) P^4^-(5′-adenosyl) tetraphosphate (Ap4A); P^1^-(5′-adenosyl) P^4^-(5′adenosyl) triphosphate (Ap3A); or P^1^-(5′-adenosyl) P^4^-(5′-guanosyl) tetraphosphate (Ap4G). Reactions were carried overnight at 37 °C in a total volume of 50 μl. PglZ (2 μM) was also incubated under the same conditions in the presence of BrxB (10 μM) and BrxB R46A (10 μM). Reactions were centrifuged at 12,000 x *g* for 10 min at 4 °C to remove precipitants and 2 μl was loaded onto an Aeris 5 μm PEPTIDE XB-C18 (150 x 4.6 mm) reversed phase high-performance liquid chromatography (HPLC) column (Phenomenex) at a flow rate of 1.5 ml/min and a linear gradient of 0-30% buffer 2 in 12 column volumes (CV), using buffer 1 (10 mM triethylammonium acetate pH 8.0) and buffer 2 (80% (v/v) acetonitrile, 10 mM triethylammonium acetate pH 8.0) in 12 CV. Protein sample in the absence of nucleotide, and nucleotide in the absence of protein sample were used as controls. Standard mixes contained 10 μM of nucleotide(s) made up to 50 μl in 1x ZB and stored at 4 °C.

### Inductively coupled plasma mass spectrometry (ICP-MS)

Total metal contents of protein samples were determined via ICP-MS (Thermo Scientific iCAP RQ ICP-MS) under KED mode (Kinetic Energy Discrimination) utilised with helium. Protein samples were diluted into 2.5% nitric acid containing 10 μg/l berylium, indium and silver as internal standards. Concentrations determined via comparison to matrix-matched elemental standard solutions.

### Analytical size exclusion chromatography

Analytical size exclusion chromatography (SEC) was performed on an ÅktaÔ Pure FPLC system (Cytiva). Protein samples were made up to 10 μM in a 100 μl final volume with analytical SEC buffer (20 mM Tris HCl pH 7.9, 150 mM NaCl). BrxB was loaded onto a Superdex™ 75 increase 10/300 GL SEC column (S-75i; Cytiva). PglZ and derivative mutants were loaded onto a Superdex™ 200 increase 10/300 GL SEC column (S200i; Cytiva). PglZ and mutant derivatives pre-incubated with BrxB and mutant derivatives at equimolar concentrations for 15 min at room temperature were also loaded onto an S200i. All columns were pre-equilibrated with 1.2 CV analytical SEC buffer. Samples were loaded onto a 100 μl capillary loop using a 100 μl Hamilton syringe. The loop was washed with 500 μl nuclease-free water followed by 500 μl analytical SEC buffer before and between each run using a 500 μl Hamilton syringe. Samples were loaded onto the column by running 500 μl of analytical SEC buffer through the capillary loop at a flow rate of 0.5 ml/min, and samples were resolved on the column using 1.2 CV analytical SEC buffer. In cases where the content of chromatogram peaks required verification by SDS-PAGE or mass spectrometry, 0.5 ml fractionation was performed, and fractions were collected in 96-well deep-plate blocks.

Calibration curves were generated by plotting the elution volumes (*V*_e_) of controls from calibration kits (GE healthcare) against their respective known molecular weights (*M*_r_). Calibration samples were prepared in 2 individual mixtures, Mix A (3 mg/ml RNase A, Ferritin, Conalbumin, Carbonic Anhydrase) and Mix B (3 mg/ml RNase A, Aldolase, Aprotinin, 4 mg/ml Ovalbumin) and made up to a final volume approximately equal to 0.5% geometric column volume. For determination of column void volume (*V*_o_), 1 mg/ml Blue Dextran was applied to the column as above, with elution volume directly proportional to *V*_o_. Elution volumes (*V*_e_) were calculated using the Peaks function in Unicorn™ 7 (Cytiva) and converted to partitioning coefficients (*K*_av_) using the following equation:

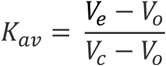

Molecular weight and Stokes radius calibration curves were subsequently plotted using Prism (GraphPad) as *K*_av_ vs Log_10_(*M*_r_,kDa) and Log_10_(*R*_st_,Å) vs *K*_av_, respectively. Observed *R*_st_ values were generated by performing linear regression on respective plots using the following equations:

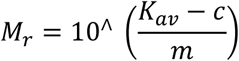

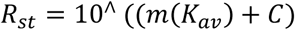

Observed values were compared against calculated hydrodynamic radii. Radius calculations of inputted AlphaFold predictive models were performed using the HullRad tool (Fluidic Analytics).

### Mass photometry

Mass Photometry experiments were undertaken on the TwoMP (Refeyn) instrument, using the Acquire 2024R1.1 and Discover 2024R1.0 software for data acquisition and analysis, respectively. DiscoverMP v2024 R2.1 was used to make figures. The autofocus function was used to find the focus plane using 19 µl of PBS on uncoated glass slides (Refeyn). Thyroglobulin monomer and dimer peaks, conalbumin and aldolase were used as standards. Stocks of each protein or complex to be tested were prepared at 100 nM immediately before dilution 1:19 into PBS and collection of a 1 min video. Gaussian fits were used for most measurements, with PgZ:BrxB and PglZ:BrxB^E74A^ measurements also making use of the interval function.

### Mass spectrometry

Collected BrxB peaks were buffer exchanged into 10 mM ammonium bicarbonate using a 10 kDa MWCO spin concentrator and submitted for positive ion electrospray time-of-flight mass spectrometry (ES^+^-ToF MS) at a final concentration of 0.5 mg/ml. Analysis was performed at our in-house Durham University Chemistry Department facility by Mr Peter Stokes using a Xevo QToF (Waters, UK) mass spectrometer.

Relevant protein bands were excised from SDS-PAGE gels for their identity to be confirmed via trypsin digest and mass spectrometry by Dr. Adrian Brown at the Department of Biosciences, Durham University.

### Pacific biosciences sequencing

Libraries for methylation sequencing were prepared using the SMRTbell HiFi 96 Prep kit (Pacific Biosciences). Bacterial gDNA was sheared using Qiagen Tissue Lyser II at 30 Hz for 240 s to produce DNA fragments with a mean size of 8–10 kb. The DNA was damage and end repaired. SMRT-bell adapters were then ligated. Exonuclease treatment removed non-incorporated SMRT-bell DNA. Sequencing was performed on a PacBio Revio (Pacific Biosciences). Data were analysed using PacBio SMRTAnalysis on SMRTLink_25.1 software Base Modification Analysis for Sequel data, to identify DNA modifications and their corresponding target motifs.

### Thermal shift assays (TSAs)

Thermal shift assays (TSAs) were performed to determine the ability of proteins to bind divalent metal cations. Samples of PglZ, PglZ incubated with 1 mM EDTA (PglZ + EDTA), and PglZ treated with EDTA (removed, as described previously; PglZ - EDTA) were incubated with 4 x 10^−3^ μl SPYRO™ Orange protein dye (ThermoFisher) per 1 μl protein for 1 h at 4 °C. Reactions containing 5 μM protein in 1x ZB were incubated with (PglZ - EDTA) or without 0.5 mM MgCl_2_, MnCl_2_, CaCl_2_, CuCl_2_, NiSO_4_, and ZnCl_2_ for 15 mins at RT, made up to 20 μl with nuclease-free water in a sealed 96-well semi-skirted PCR plate (Starlab). Samples were centrifuged and inserted into a CFX connect real-time qPCR machine for thermal shift analysis. The fluorescence was measured in 0.5 °C increments from 25 to 95 °C. Deconvolution of thermal shift isotherms was performed using NAMI python tool (28), and thermal shift graphs were generated using Prism (GraphPad).

### Nuclease assays

The ability of PglZ to degrade DNA and RNA was analysed using plasmid DNA, phage gDNA, and phage RNA. Prior to any assays, PglZ was treated with EDTA, as described previously, to ensure consistency in the metal binding between samples. For titration experiments, 0, 12, 24, 48, 96, 192, 384, 768, and 1536 nM of purified PglZ were incubated with 6 nM pSG483 supercoiled plasmid DNA, 6 nM pSG483^BREX^ ^KO^ supercoiled plasmid DNA (*E. fergusonii* BREX site mutated), 200 ng pBrxXL WT plasmid DNA, and 200 ng pBrxXL-Δ*pglX* plasmid DNA. Purified PglZ at 0, 48, 96, 192, 384, 768, and 1536 nM was also incubated with 200 ng φT4 gDNA, 200 ng φPau gDNA, 6 nM M18mp13 ssDNA, and 6 nM φMS2 RNA. Reactions were incubated for 1 h at 37 °C with 1x ZB and 0.5 mM MnCl_2_. Control reactions either eliminated the metal or included 1 mM ATP in the reaction mix. The activity of PglZ derived mutants against supercoiled pSG483 were tested at 384, 768, and 1536 nM in the presence of 1x ZB and 0.5 mM MnCl_2_ at 37 °C for 1 h.

To test the activity of PglZ in the presence of various metals, PglZ (768 nM) was incubated with supercoiled pSG483 (6 nM) in 1x ZB at 37 °C for 1 h in the presence of 0.5 mM MgCl_2_, MnCl_2_, CaCl_2_, ZnCl_2_, CuCl_2_, and NiSO_4_. Control reactions contained no divalent cations. To test the inhibition of PglZ by various nucleotides, PglZ (768 nM) was incubated supercoiled pSG483 (6 nM) in 1x ZB and 0.5 mM MnCl_2_ with and without 1 mM ATP, GTP, CTP, UTP, dATP, dGTP, dTTP, dCTP, ADP, AMP, or AMP-PNP for 1 h at 37 °C. Control reactions contained no nucleotide or no PglZ.

To test the activity of PglZ in the presence of BrxB, PglZ (768 nM) was incubated with pSG483 (6 nM) with BrxB at 0.35, 0.7, 1.5, 3.0, and 6.1 μM for 1 h at 37 °C. Reactions comprised of 1x ZB and 0.5 mM MnCl_2_ and were completed in the presence and absence of 1 mM ATP. PglZ was also incubated with BrxB mutants W135A and R46A. Control reactions contained no protein, PglZ only (768 nM), and BrxB or derivative mutants only (6.1 μM).

All reactions were made up to 20 μl. Reactions were stopped by the addition of 2 μl stopping buffer (5% SDS (v/v), 125 mM EDTA) followed by 4 μl TriTrack loading dye (ThermoFisher). Samples were analysed by agarose gel electrophoresis in 1.4% (w/v) gels for pSG483 analysis, or 0.8% (w/v) gels for pBrxXL, phage gDNA, and RNA analysis using 1x TAE buffer and running at 45 V for ∼16 h. Agarose gels were post stained in 1x TAE containing 0.5 μg/ml ethidium bromide and destained in 1x TAE. Gel images were obtained on a ChemiDoc™ Imaging System on the ethidium bromide setting (BioRad). Gel images were analysed using Fiji (ImageJ; v 2.1.0) with background subtracted. For pSG483 assays, the supercoiled, nicked, and linear DNA band intensity was measured per lane and calculated as a percentage of the total DNA in the respective lane. For pBrxXL assays, the DNA band intensity of all bands per lane were measured independently and compared as a percentage against the corresponding DNA band in the ‘0’ PglZ control lane. For phage gDNA/RNA assays, intact phage gDNA/RNA band intensity was measured in each lane and compared as a percentage against the ‘0’ PglZ control. Mean values and standard deviation were calculated from triplicate data. Data were plotted in Prism (GraphPad).

### Efficiency of plating (EOP)

*E. coli* bacteriophages were isolated from freshwater sources in Durham city centre and the surrounding areas, as described previously (23). *E. coli* DH5α were transformed with pTRB563 (pBrxXL), pTRB564 (pBrxXL-Δ*pglX*), pTRB744 (pBrxXL-*brxB* W135A), pTRB745 (pBrxXL-*pglZ* H741A), pTRB746 (pBrxXL-*brxB* E47A), pTRB747 (pBrxXL-*brxB* E89A), pTRB748 (pBrxXL-*brxB* S133A), pTRB749 (pBrxXL-*brxB* R46A), pTRB750 (pBrxXL-*pglZ* T538A), or pTRB766 (pBrxXL-*pglZ* T538A/H741A) and grown overnight. Serial dilutions of phage Pau were produced in phage buffer (10 mM Tris HCl pH 7.4, 10 mM MgSO_4_, 0.01% (v/v) gelatin). 200 μl of overnight culture and 10 μl of phage dilution were added to a sterile 8 ml plastic bijoux with 3 ml of 0.35% (w/v) LB-agar and poured onto LB plates. Plates were incubated overnight at 37 °C before plaque forming units (pfu) were counted on each plate. EOP values were calculated by determining the phage titre on a test strain divided by the titre on a control strain. EOP data were collected in triplicate and the mean value was plotted in GraphPad Prism.

### Initial single particle screening of *Salmonella* PglZ:BrxB

Negative stain grids were prepared by applying 4 μl of size exclusion chromatography (SEC) purified PglZ:BrxB sample at a concentration of approximately 0.04 mg/ml to a glow-discharged Formvar/Carbon 400 mesh Copper grid (Ted Pella). The sample was allowed to absorb for 30 s followed by wicking excess solution with filter paper. The grid was quickly washed two times in 30 μl drops of water and once in a 30 μl drop of 2% uranyl formate (UF) followed by a final staining for 30 s with another 30 μl drop of 2% UF. The grids were air dried for at least 1 hr. Grids were screened on an in-house Talos L120C transmission electron microscope (Thermo Fisher), operating at 120 kV and equipped with a 4k x 4k Ceta CMOS high-resolution 16M camera (Thermo Fisher). The sample distributed homogeneously and at random orientations over the surface of the prepared negative stained grids.

### CryoEM hybrid model determination for *Salmonella* PglZ:BrxB

Flow charts and summary of data collection of the methods described below are shown in **Supplementary Figure S2**. Grids were prepared for cryoEM by applying 3 μl of SEC purified sample at a concentration of 0.25 or 0.5 mg/ml (diluted in 20 mM Tris pH 8.0, 300 mM KCl) to a glow-discharged C-Flat 1.2/1.3 holey carbon film coated copper grid (Electron Microscopy Sciences). The grids were blotted for 5 s at a tension of 0, and plunge-frozen into liquid ethane using a Mark IV Vitrobot (Thermo Fisher). Two datasets of 4686 (dataset 1) and 4731 (dataset 2) movies were collected at a super resolution pixel size of 0.56 Å using a Glacios 200 kV electron microscope (Thermo Fisher) equipped with a Gatan K3 direct electron detector. Preprocessing of datasets was performed in WARP (29) where pixels were binned to 1.122 Å. Datasets were imported into CryoSPARC (30) and particles in dataset 1 were picked by automated searching for Gaussian signals, extracted and Fourier cropped to a box size of 300 and 100 pixels, respectively, and filtered with multiple rounds of 2D classification and selection. Final particles from dataset 1 were lowpass filtered to 20 Å and used as a template for particle picking in dataset 2. Picked particles from dataset 2 were then extracted and filtered as in dataset 1. Final particles from both datasets were combined into a single Ab-initio 3D reconstruction job with 4 classes, resulting in a single class with full particles (124,721) and the remaining classes with fragments or junk particles. The particles contained in the single class were reextracted without Fourier cropping to a box size of 300 pixels followed by homogeneous and non-uniform refinement (31) resulting in a map with GSFSC resolution of 4.45 Å. The resulting volumes were evaluated in ChimeraX (32).

### Model fitting

Predicted models of the *Salmonella* PglZ:BrxB sub-complex was generated by AlphaFold (33) resulting in high per-atom confidences. Initial placement of models was accomplished in ChimeraX (32) using the Fit in Map tool. Domains were then further fit into the volume individually. The predicted PglZ:BrxB interface was preserved by treating PglZ residues 1-98 as part of the BrxB domain. The models were then further refined in the Phenix suite (34) using Cryo_fit (35) and Real Space Refine (36). No rebuilding was performed due to lack of detail in the volumes.

## Results

### Salmonella *BREX components form larger complexes* in vivo

Having previously performed characterisation of independent core Type I BREX components BrxA, PglX and BrxL (17, 18, 21, 22), we turned to examining interactions between BREX proteins. The BREX locus from *Salmonella* Typhimurium strain D23580 (**Fig. 1A**) had already been sub-cloned and shown to be active in phage defence (17, 23). *Salmonella* genes *brxA*, *brxB, brxC* and *pglX* were cloned into one multiple cloning site of pCOLA DUET1, and genes *pglZ* and *brxL* were cloned into the second multiple cloning site. A compatible vector based on 2HR-T (addgene #29718) was generated that expressed His-strep-BrxB. Combining these two vectors, and appropriate vector-only controls, we observed robust expression of the His-strep-BrxB fusion in the absence of the full BREX locus (**Fig. 1B**). When then expressed in cells also expressing the full *Salmonella* BREX locus we observed co-purification of BREX proteins BrxC, PglX, PglZ and BrxL with His-Strep-BrxB, indicating formation of higher order complexes (**Fig. 1B**). The indicated bands were confirmed for identity by mass spectrometry (**Fig. 1B**). The most abundant protein after His-Strep-BrxB was PglZ. In order to produce larger quantities of the BREX complex(es) the six BREX genes were cloned as three pairs into compatible DUET vectors and co-purification was performed on strains containing increasing combinations of expression vectors (**Fig. 1C**). For these experiments, the His-strep tag was replaced with a His-SUMO tag to aid later purification. We saw robust His-SUMO-BrxB co-purification with PglZ, and then with PglZ, BrxC and PglX, and finally again PglZ, BrxC, PglX and BrxL. No BrxA was co-purified (**Fig. 1C**). We noted that certain combinations of vector caused poor growth of cells and so performed viable counts (**Fig. 1D and Supplementary Fig. S1A**). Expression of BrxC and PglX was toxic in *E. coli*, but this was in part negated by co-expression of His-SUMO-BrxB and PglZ, or BrxA and BrxL, or all six proteins. Due to the robust expression and co-purification of His-SUMO-BrxB and PglZ we chose to pursue this sub-complex for further study. Large scale co-expression and co-purificaiton of His-SUMO-BrxB and PglZ yielded a clean sample of native PglZ:BrxB sub-complexes (**Supplementary Figs. S1B-C**).

Having identified PglZ:BrxB as a strong pairwise protein-protein interaction in *Salmonella*, we further tested this observation using a previously characterised Type I BREX system found in *Acinetobacter* (20). Using that system, it was also found that PglZ and BrxB interact strongly and co-eluted from affinity-based and size exclusion columns in a 1:1 ratio (**Supplementary Fig. S1**). Interestingly, and unlike its behaviour in *Salmonella*, BrxB from *Acinetobacter* was found to require the co-expression and corresponding presence of bound PglZ in order to remain soluble *in vitro.* These data indicate that the PglZ:BrxB interaction is generalisable and reproducible.

### Salmonella *PglZ:BrxB complexes show dynamic movement*

Size exclusion chromatography (SEC) of native PglZ, BrxB and co-expressed and co-purified PglZ:BrxB expected sub-complexes demonstrated an altered elution profile for PglZ:BrxB, indicating formation of a larger sub-complex (**Fig. 2A**). The *Salmonella* PglZ:BrxB sub-complexes were used to perform structural studies through negative stain transmission electron microscopy, followed by cryo-electron microscopy (cryoEM) (**Fig. 2B and Supplementary Fig. S2**). The final model had a Gold Standard Fourier Shell Correlation (GSFSC) resolution of 4.45 Å (**Supplementary Fig. S2**). At this resolution it was possible to make use of AlphaFold outputs for PglZ:BrxB to generate a final hybrid model of the PglZ:BrxB sub-complex. BrxB is itself a globular protein and was shown to be bound to the N-terminal domain (residues 1-96) of PglZ (**Fig. 2B**). EMBL PISA (37) identified BrxB residues R49, N135 and W137, and PglZ residues K58, E62 and D88, as important for binding (**Fig. 2B, inset**). PglZ forms an “S” shape, with a central domain (residues 98-292) and a large C-terminal PglZ domain (residues 304-748) that contains the metal-binding site identified within PorX (16), and a final extension including a seven sheet β-barrel (residues 749-867) (**Fig. 2B**). Comparison of the *Salmonella* PglZ:BrxB AlphaFold model alone against our cryoEM hybrid model shows two distinct points of movement (**Supplementary Fig. S3**). The PglZ N-terminal domain and BrxB, and the C-terminal β-barrel extension have both made large movements between the two models, whereas the central and PglZ domains remain fixed (**Supplementary Fig. S3**). This dynamic flexibility is likely the cause of our data being limited to lower resolution. Nevertheless, this model confirms the presence of a flexible but stable PglZ:BrxB complex.

**Figure 2.**
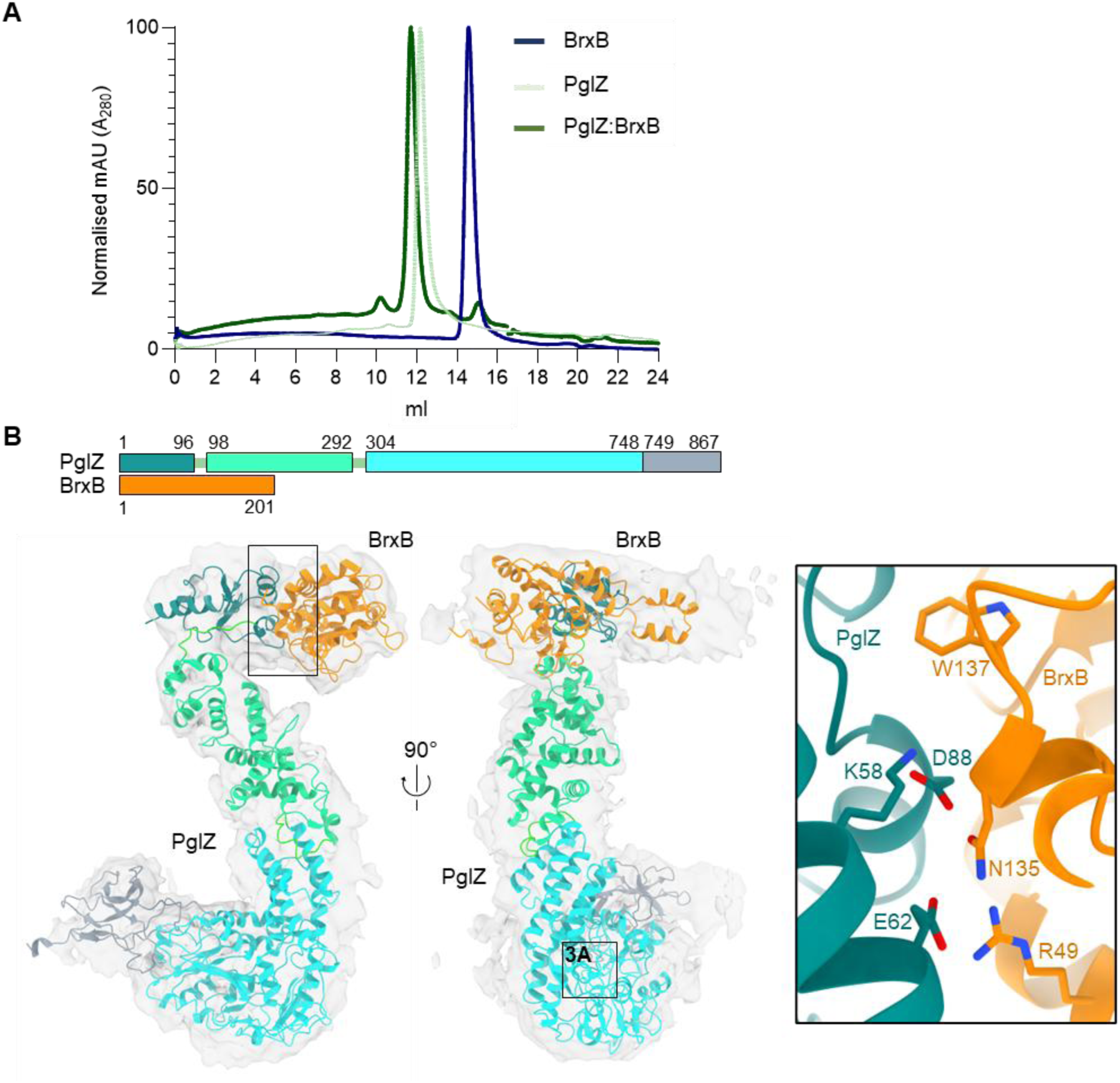
Structure of the *Salmonella* PglZ:BrxB stable sub-complex. (**A**) Size exclusion chromatography traces of independent *Salmonella* PglZ and BrxB purifications, and the PglZ:BrxB co-expression sample used for cryoEM, show that PglZ and BrxB form a stable complex. (**B**) A cryoEM map generated from single particles of the *Salmonella* PglZ:BrxB complex validates the interaction between both proteins, as well as the location of the PglZ:BrxB interaction surfaces and identity of residues in the protein-protein interface (inset box). The resulting model is closely related to the corresponding AlphaFold prediction for the *Salmonella* PglZ:BrxB complex, albeit with slight rearrangements corresponding to a small rotation of the N-terminal domain of PglZ and associated BrxB relative to the larger core of PglZ (**Supplementary Figure S3**).

### PglZ can cleave cyclic and linear oligonucleotides in a metal-dependent manner

Next, we performed biochemical characterisation of PglZ in isolation, in preparation for later investigation of the PglZ:BrxB sub-complex. As experimentation began we noted that the *Salmonella* PglZ homologue had a tendency to precipitate during tests. As such, we chose to use PglZ from *E. fergusonii*, a system we had previously characterised (24), as a substitute biochemical model.

A superposition of the AlphaFold output for the PglZ domain from *E. fergusonii* PglZ (residues V474-L759) with the PglZ domain from PorX (PDB: 7PVK, residues 213-518) produced an RMSD of 2.822 (over 1096 atoms) (**Figs. 2B and 3A**). Residues identified as important for PorX metal binding and oligonucleotide cleavage activity, T272 (mutated to T272A in PDB 7PVK) and H500 (16) correspond to *E. fergusonii* PglZ residues T538 and H741, respectively (**Fig. 3A**). Mutant proteins *E. fergusonii* PglZ T538A, H741A and a double mutant T538A/H741A were expressed and purified, and shown to have similar mass photometric and SEC profiles as *E. fergusonii* PglZ WT (**Supplementary Fig. S4**).

**Figure 3.**
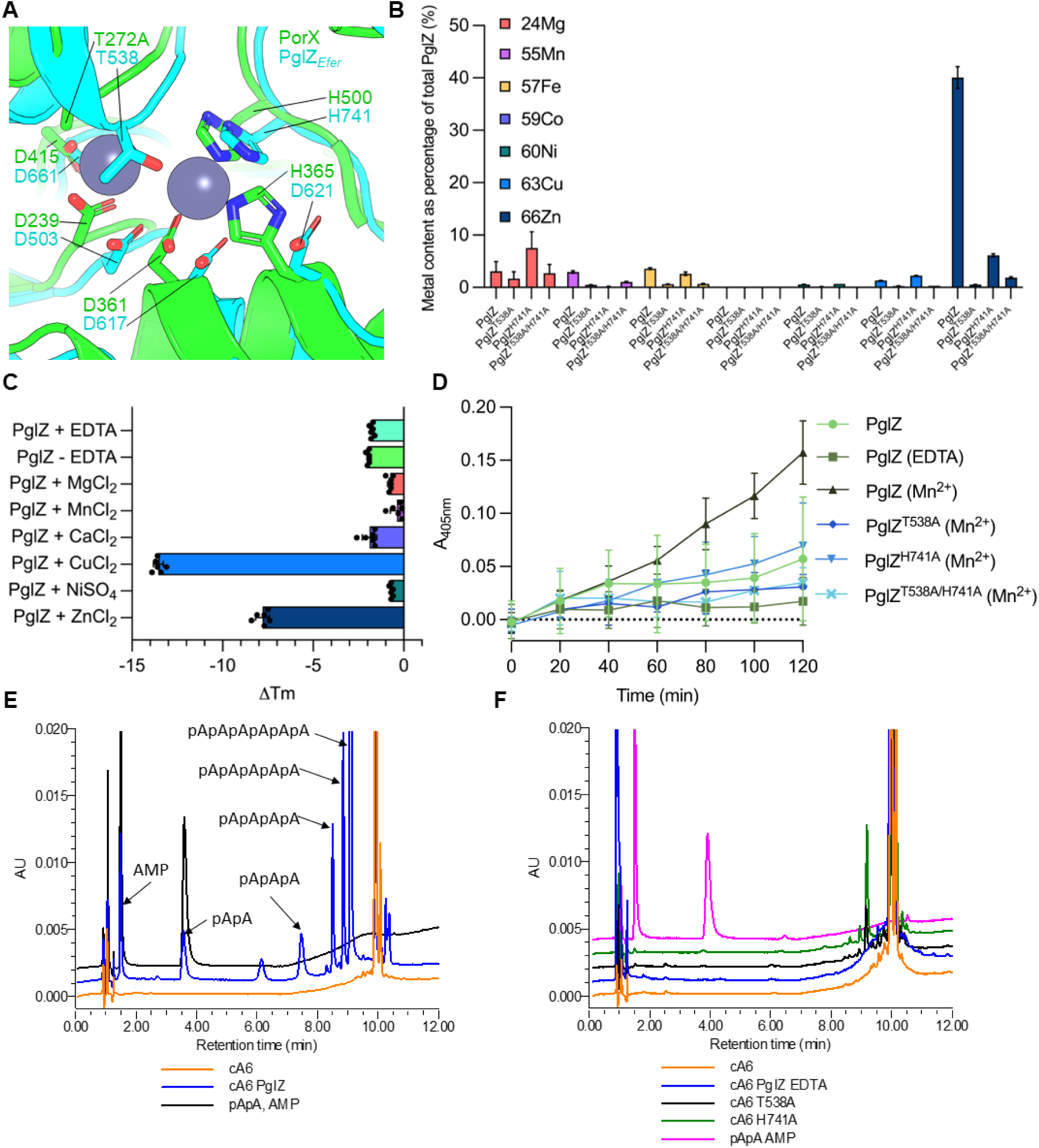
PglZ can cleave cyclic nucleotides in a metal-dependent manner. (**A**) Overlay of AlphaFold2 *E. fergusonii* PglZ predicted structure with PorX from *Porphyromonas gingivalis* (PDB: 7PVK). (**B**) ICP-MS of PglZ WT and mutants showing divalent cations bound following purification. Plotted data represent mean values ± SD. Metal content is plotted as a percentage of the total protein in the sample. (C) Thermal shift assays performed upon PglZ WT (5 μM) following incubation with EDTA or metals (0.5 mM). Mean changes in melting temperature (ΔTm) are plotted by comparison to PglZ WT in the absence of EDTA or metal, which is set as ‘0’. Error bars represent standard deviation (6 replicates) (D) Bis(4-nitrophenyl) phosphate (2.5 mM) phosphodiesterase assays using PglZ WT (2 μM) in the presence and absence of EDTA and Mn, and mutants in the presence of Mn. The Mn is supplied by 0.5 mM MnCl_2_. Plotted data represent mean values ± SD (9 replicates). Absorbance (A_405nm_) represents the amount of reaction product *p*-nitrophenyl phosphate. (**E**) HPLC analysis of cyclic hexa-adenosine monophosphate (cA6) cleavage by PglZ WT (2 μM) in the presence of Zn (10 μM). (**F**) HPLC analysis of cyclic hexa-adenosine monophosphate (cA6) cleavage by PglZ WT (2 μM) treated with EDTA, and mutants T538A and H741A in the presence of Zn (10 μM). Control reactions are represented by cA6 alone, and standard mixes are comprised of pApA and AMP. All nucleotides in the reaction mixes are at 10 μM. Presented traces are representative of triplicate data.

PglZ wild type (WT) and mutants were tested for metal content following purification, using inductively coupled plasma mass spectrometry (ICP-MS). There was a clear abundance of zinc in PglZ WT samples, and levels were greatly lowered in the mutant samples (**Fig. 3B**). Following EDTA treatment to remove metals and subsequent purification to remove EDTA, *E. fergusonii* PglZ WT and mutants were tested for stability in the presence of a range of metals using thermal shift assays (TSAs) (**Fig. 3C and Supplementary Fig. S5**). PglZ proteins were destabilised by copper and, surprisingly, zinc, but were stabilised by magnesium, manganese and nickel (**Fig. 3C and Supplementary Fig. S5**). Calcium had no effect, likely because it could not bind (**Fig. 3C**). Mutants PglZ T538A and PglZ H741A were stabilised by manganese, indicating some metal binding could occur (**Supplementary Fig. S5**). Double mutant PglZ T538A/H741A was not stabilised by any metal indicating that metal binding in the active site was no longer possible (**Supplementary Fig. S5**). The double mutant was, however, still destabilised by copper and zinc, suggesting effects for copper and zinc seen with both this mutant and also PglZ WT are due to non-specific binding (**Supplementary Fig. S5**). The melting temperatures for PglZ WT and mutants indicated that T538A reduces overall stability, but H741A has less of an impact (**Supplementary Fig. S5E**).

Initial tests for potential phosphodiesterase activity using bis-*p*NPP as a substrate with PglZ WT and additional zinc resulted in precipitation, and so magnesium, manganese, and calcium were all tested as alternates, with manganese showing the greatest levels of activity (**Supplementary Fig. S6**). Manganese was therefore selected as an alternate metal in bis-*p*NPP phosphodiesterase activity assays. Having stripped metals from the samples and restored manganese, *E. fergusonii* PglZ WT showed robust production of *p*-nitrophenol from bis-*p*NPP, at levels greater than for untreated PglZ WT that contained the zinc remaining after purification (**Fig. 3D**). In contrast, when all mutants were EDTA treated, re-purified, and then provided manganese, PglZ H741A had reduced activity, and both PglZ T538A and the double mutant T538A/H741A lacked activity, demonstrating levels similar to those observed for the PglZ WT sample that was without metal following EDTA treatment (**Fig. 3D**). This result indicated that PglZ, like PorX, can cleave bis-*p*NPP in a metal-dependent manner, and that mutations interfering with the likely metal binding site caused reduced activity.

Cleavage of cyclic oligonucleotides was then tested by incubating *E. fergusonii* PglZ WT with cA6 and analysing the resulting products by high performance liquid chromatography (HPLC). Having tested a range of metals it was noted that zinc was the preferred metal in these assays, and was used at a reduced concentration to prevent protein destabilisation and precipitation. *E. fergusonii* PglZ WT robustly linearised cA6 and sequentially cleaved nucleotide products, indicated by a trace for each linear species down to AMP (**Fig. 3E**). PglZ cleavage activity was ablated by EDTA treatment, and mutant PglZ T538A showed no appreciable activity (**Fig. 3F**). Mutant H741A appeared able to cleave cA6 but did not produce further cleavage products (**Fig. 3F**). PglZ was then tested against a broader range of nucleotides (**Supplementary Figs. S7 and S8**). PglZ was observed to cleave cyclic oligonucleotides (cA4, cA3, cA2, and cG2) and linear oligonucleotides (pApA, pApG, pGpG, c(ApGp), and c[A(3′5′)pA(3′5′)pG(3′5′)]) containing both adenosine and guanosine, including an oligonucleotide with 2′-5′ rather than 3′-5′ phosphodiester linkages (c[G(2′5′)pA(3′5′)p]). PglZ was unable to cleave cyclic mononucleotides (cAMP, cGMP, cTMP, cUMP, cCMP, 2′3′ cAMP, 2′3′ cGMP, and 2′3′ cUMP) or dinucleotide polyphosphates (Ap3A, Ap4A, and Ap4G) in the presence of Zn. Following this analysis we returned to testing metal usage and noted that at low manganese concentrations we were also able to observe PglZ-dependent cleavage of both cA6 and pApA (**Supplementary Fig. S7C**). Collectively, these data show robust metal-dependent cyclic and linear oligonucleotide cleavage by PglZ from a BREX system.

### PglZ is an endonuclease that can cleave dsDNA

Having established that PglZ has nuclease activity against oligonucleotides we were curious as to whether PglZ could cleave dsDNA. Plasmid pSG483, a pUC19 derivative that can be easily prepared as supercoiled (S), nicked (N), relaxed (R) or linear (L) dsDNA, was selected as a suitable substrate to be tested against *E. fergusonii* PglZ. Initial assays indicated that manganese would be the preferred metal in this context, but supplementing with zinc did allow some PglZ nuclease activity (**Supplementary Fig. S9A**). Incubation of pSG483 with a titration of *E. fergusonii* PglZ WT revealed both nicking and linearisation activities, which were metal-dependent and could be inhibited by ATP (**Fig. 4A**). This confirmed that PglZ can nick and cut dsDNA, and is an endonuclease. Due to the observed inhibition by ATP, a range of mononucleotides were then tested. All NTPs, dNTPs and AMP-PNP inhibited PglZ nuclease activity, but AMP did not (**Supplementary Fig. S9B**). The PglZ mutants were then tested for nuclease activity (**Fig. 4B**). *E. fergusonii* PglZ H741A had increased nicking but decreased linearisation activity, whereas PglZ T538A and the double mutant PglZ T538A/H741A were both ablated for activity (**Fig. 4B**). This follows the previous observed trend for activity (**Fig. 3F**) and indicates T538A prevents metal binding and therefore activity, whilst H741A reduces metal binding and activity.

**Figure 4.**
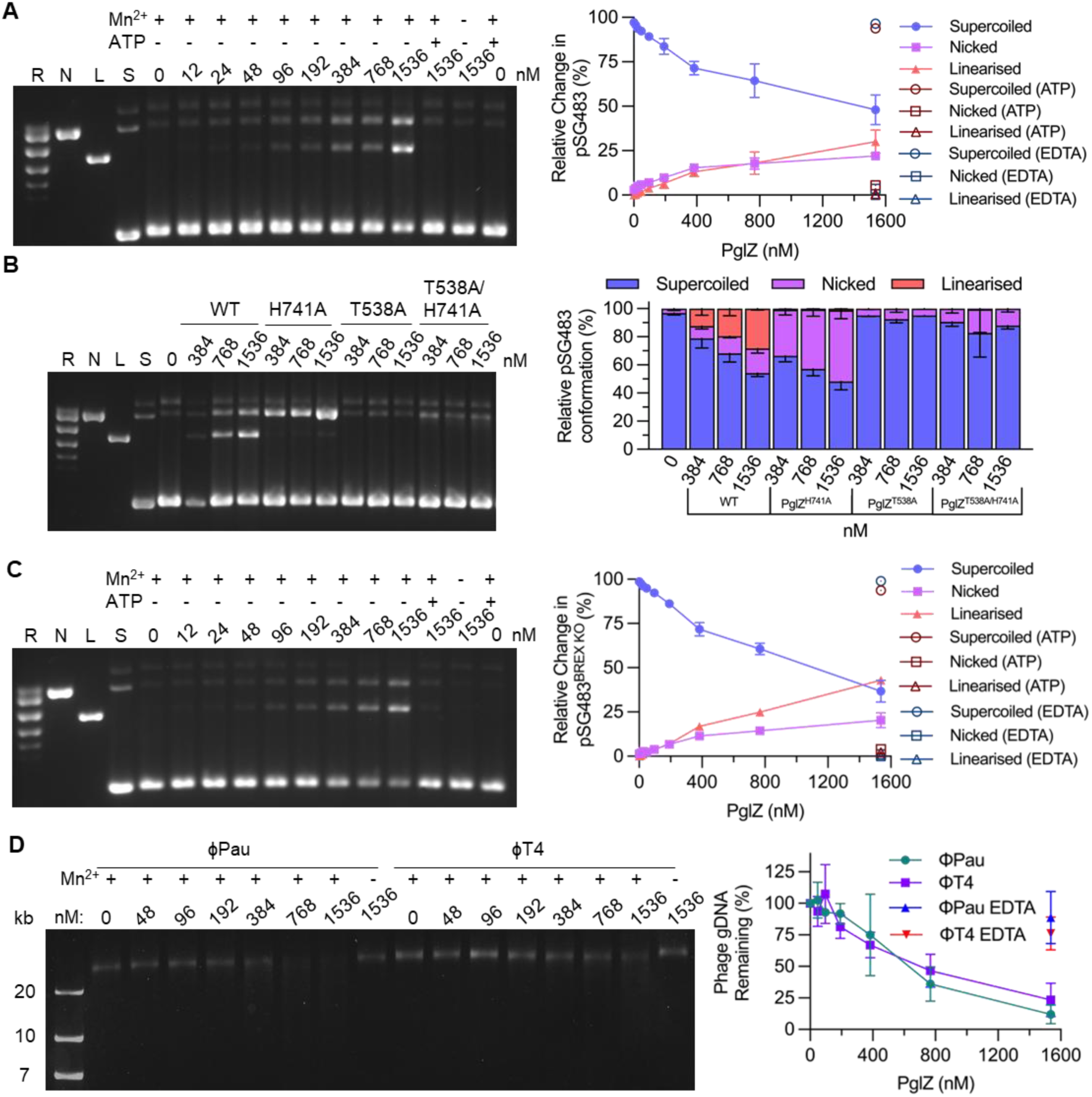
PglZ is a metal-dependent nuclease that does not recognise BREX sites. (**A**) PglZ nicks and linearises supercoiled plasmid pSG483 in a metal-dependent manner that is inhibited by ATP. (**B**) Mutation of metal binding site alters nuclease activity, either by shifting activity away from linearisation and towards nicking (H741A) or eliminating activity (T538A and T538A/H741A). (**C**) PglZ can linearise plasmid pSG483 DNA with a mutated BREX site. (**D**) PglZ does not appear to have site-specific nicking activity on linearised phage DNA, nor be impacted by DNA modifications such as those found in T4. PglZ was titrated against constant supercoiled pSG483 plasmid DNA (6 nM) or phage gDNA (200 ng) in the presence and absence of MnCl_2_ (0.5 mM) and ATP (1 mM). Control lanes represent supercoiled (S), nicked (N), linear (L), and relaxed (multiple topoisomers; R) plasmid DNA, or contain the appropriate DNA in the absence of protein. Assays are presented on 1.4% (w/v) or 0.8% (w/v) agarose 1x TAE gels post-stained with ethidium bromide. Assays shown are representative of triplicate experiments. Data points and error bars represent the mean ± SD of triplicate data.

The BREX methyltransferase, PglX, determines sequence recognition for host methylation and phage defence (17). *E. fergusonii* PglX recognises the sequence GCTAAT, and there is 1 copy of this motif in pSG483. A mutant pSG483 was generated (pSG483^BREX^ ^KO^) with the GCTAAT motif mutated to GCTATT to allow testing of whether PglZ cleavage is dependent on BREX motifs. When a titration of *E. fergusonii* PglZ WT was titrated against pSG483^BREX^ ^KO^ there was no observed difference to the result with pSG483 (**Figs. 4A and 4C**). We then considered whether BREX methylation might impact PglZ nuclease activity. We prepared pBREXxl WT, a plasmid encoding the full *E. fergusonii* locus and the mutant pBREXxl-Δ*pglX*. Each plasmid has previously been shown to be BREX methylated and lacking methylation, respectively (24). *E. fergusonii* PglZ WT caused equal degradation of both plasmids, indicating BREX methylation does not impact PglZ activity under these isolated conditions (**Supplementary Figs. S9C-D**).

Phage Pau was shown to be susceptible to *E. fergusonii* BREX in an earlier study (24). Phage Pau does not have modified DNA (23), unlike phage T4, which has modified cytosines and so is inherently resistant to BREX. When tested, *E. fergusonii* PglZ was able to cleave genomic DNAs from both these phages (**Fig. 4D**). The cleavage did not produce a distinct pattern, rather a faint smear of products, demonstrating that PglZ is likely a sequence-independent endonuclease (**Fig. 4D**). It was also interesting that T4 cytosine modifications did not impact PglZ cleavage when in isolation, though T4 (and other modified phages) are resistant to BREX phage defence. *E. fergusonii* PglZ WT could also cause sequence-independent cleavage of ssDNA, using M18mp13 genomic DNA as substrate (**Supplementary Fig. S10A**). There was no activity, however, on MS2 phage genomic RNA (**Supplementary Fig. S10B**).

Finally, to ensure our biochemical data and structural study are aligned, we confirmed that *Salmonella* PglZ WT also demonstrated metal-dependent nicking and linearisation of pSG483 (**Supplementary Fig. S10C**). *Salmonella* PglZ showed a preference for zinc or magnesium and in contrast to *E. fergusonii* PglZ, *Salmonella* PglZ could also use calcium and copper, and could not use manganese (**Supplementary Fig. S10C**).

### PglZ:BrxB interactions can be ablated by interface mutations

As both *E. fergusonii* and *Salmonella* PglZ were shown to be nucleases, we also wanted to demonstrate that *E. fergusonii* PglZ and BrxB also form sub-complexes as observed for the *Salmonella* and *Acinetobacter* homologues (**Figs. 1 and 2, Supplementary Fig. S1**). Our hybrid model identified several *Salmonella* BrxB residues important for the PglZ:BrxB interaction (**Fig. 2B**). A suite of *E. fergusonii* BrxB WT and mutant proteins were expressed and purified (**Supplementary Fig. S11A**). None of the proteins contained any metals after purification, as analysed by ICP-MS (**Supplementary Fig. S11B**). SEC analysis of *E. fergusonii* BrxB WT showed that it formed both monomer and dimer peaks (**Supplementary Figs. S11C and D**), as confirmed by native mass spectrometric analysis (**Supplementary Fig. S11E**). Analytical SEC was performed using *E. fergusonii* BrxB WT, PglZ and BrxB WT with PglZ (**Fig. 5A**). Incubating BrxB WT with PglZ caused higher order complexes to form, as shown by elution profiles and corresponding SDS-PAGE analysis of the peaks (**Fig. 5A**). This indicated that *E. fergusonii* PglZ:BrxB sub-complexes were also forming, though perhaps with higher order forms being produced beyond those observed for *Salmonella* homologues (**Fig. 2A**).

**Figure 5.**
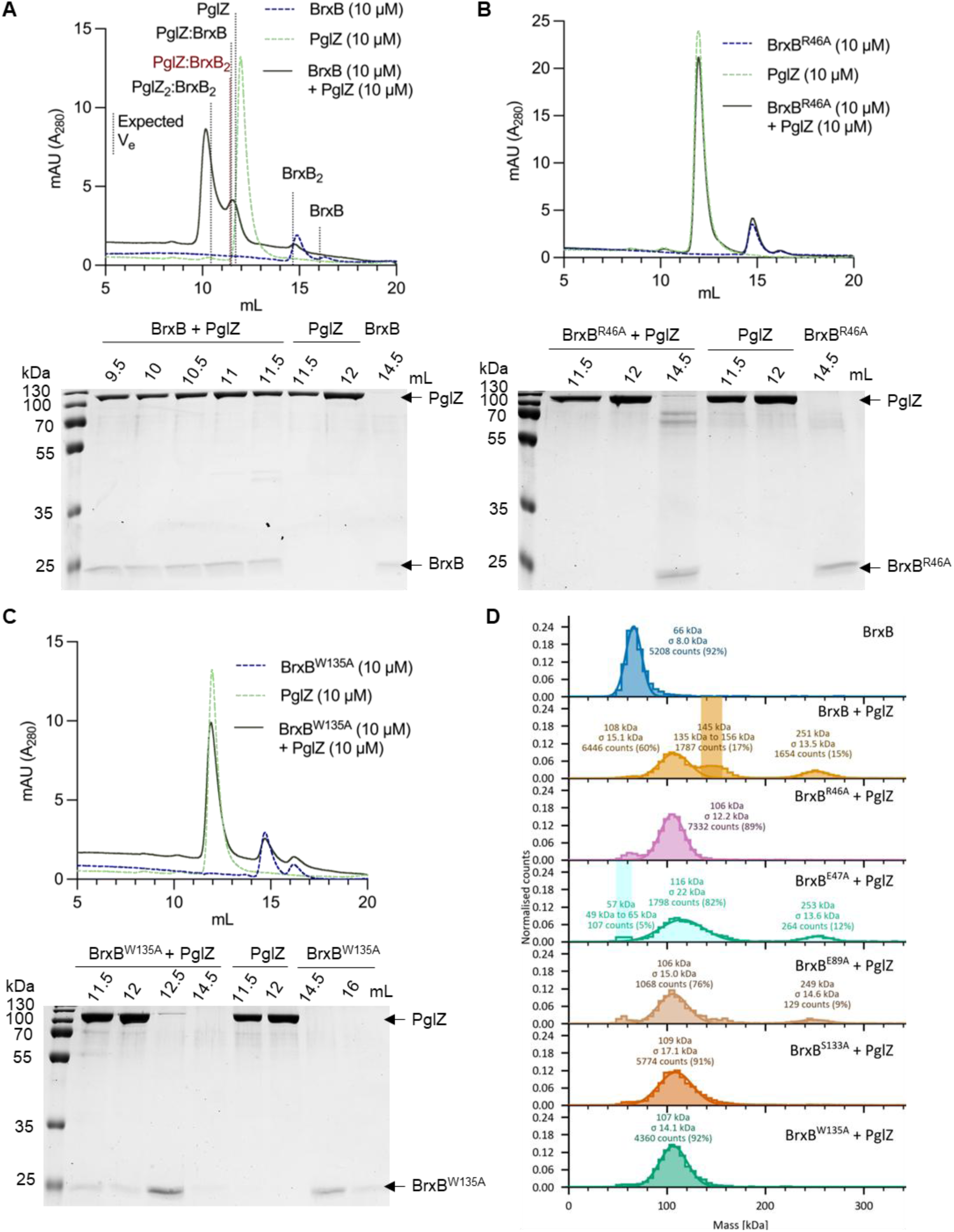
*E. fergusonii* BrxB mutants fail to form complexes with PglZ. Analytical SEC (S200i) and SDS-PAGE analysis of PglZ:BrxB complexes formed with BrxB WT (**A**), BrxB^R46A^ (**B**), and BrxB^W135A^ (**C**). Samples of PglZ (10 μM) with equimolar BrxB WT and mutants were made up to 100 μl and pre-incubated for 15 min prior to loading on the S200i. The expected elution volumes (V_e_) of various complex conformations are highlighted by black or red dotted lines, and the elution profile of PglZ incubated with BrxB is shown as a dark green solid line. Control elution profiles of PglZ alone (light green dashed line) and BrxB alone (dark blue dashed line) are also shown. Fractionated peak samples were resolved on 15% (v/v) polyacrylamide gels for 1 h 15 min in tris-glycine running buffer and stained with Quick Coomassie. Protein identities are highlighted with black arrows. (**D**) Mass photometry of PglZ with BrxB WT and mutants. Counts were acquired for 60 s with BrxB (5 nM) or samples of PglZ (5 nM) pre-incubated with equimolar BrxB WT and mutants in phosphate buffered saline (PBS).

In contrast, co-incubation of *E. fergusonii* PglZ WT with BrxB R46A and BrxB W135A failed to produce PglZ:BrxB complexes (**Figs. 5B and C**). BrxB mutants E47A, E89A and S133A generated intermediate elution profiles, indicating some complexes were forming, but to lesser extent than with BrxB WT (**Supplementary Fig. S12**). Mass photometric analysis of *E. fergusonii* PglZ incubated with BrxB WT and mutants showed similar trends, in that complexes formed with BrxB WT, none formed with BrxB R46A or BrxB W135A, or BrxB S133A in these conditions, and complexes formed but less robustly with BrxB E89A and BrxB E47A (**Fig. 5D**).

Finally, we also examined by analytical SEC whether any of the *E. fergusonii* PglZ mutants T538A, H741A or T538A/H741A would impact BrxB WT binding and formation of higher order complexes. As expected, due to these mutations being distant from the BrxB binding site (**Figs. 2B and 3A**), no impact on complex formation was observed (**Supplementary Figure S13**). Together, these data support comparisons between our two model homologues, as both were shown to have nuclease activity, and also now both have been shown to form sub-complexes.

### Interaction with BrxB impacts neither PglZ nuclease activity nor inhibition of nuclease activity by ATP

Gel-based nuclease activities were then used to investigate the impact of BrxB interactions on PglZ activity. In these assays, *E. fergusonii* BrxB had no identifiable nicking or linearisation activity (**Fig. 6A**). Titration of *E. fergusonii* BrxB against *E. fergusonii* PglZ had no appreciable impact on PglZ nicking and linearisation activities until the highest BrxB concentration (**Fig. 6A**). Comparisons of the AlphaFold model for the BrxB structure using DALI (38) indicated some similarity to nucleotide binding regions of AAA+ proteins, but BrxB appears to be lacking key Walker motif residues. As BrxB had the potential for binding ATP, we investigated whether BrxB might impact the observed inhibition of PglZ activity by ATP (**Fig. 4A**). When the same PglZ to BrxB titration was performed in the presence of ATP there was no indication that BrxB could overcome inhibition of PglZ activity by ATP (**Fig. 6A**). For completeness, we also tested whether non-interacting *E. fergusonii* BrxB mutants impacted PglZ activity, but no effect was observed (**Fig. 6B**). Next, we used HPLC analysis of oligonucleotide cleavage as another measure of BrxB impact. Neither BrxB WT nor BrxB R46A altered the ability of *E. fergusonii* PglZ to cleave cA6 or pApA (**Fig. 6C**). This indicates that the role of BrxB, at least in these isolated conditions, is independent of PglZ nuclease activity, and likely has more relevance in the context of larger BREX complexes.

**Figure 6.**
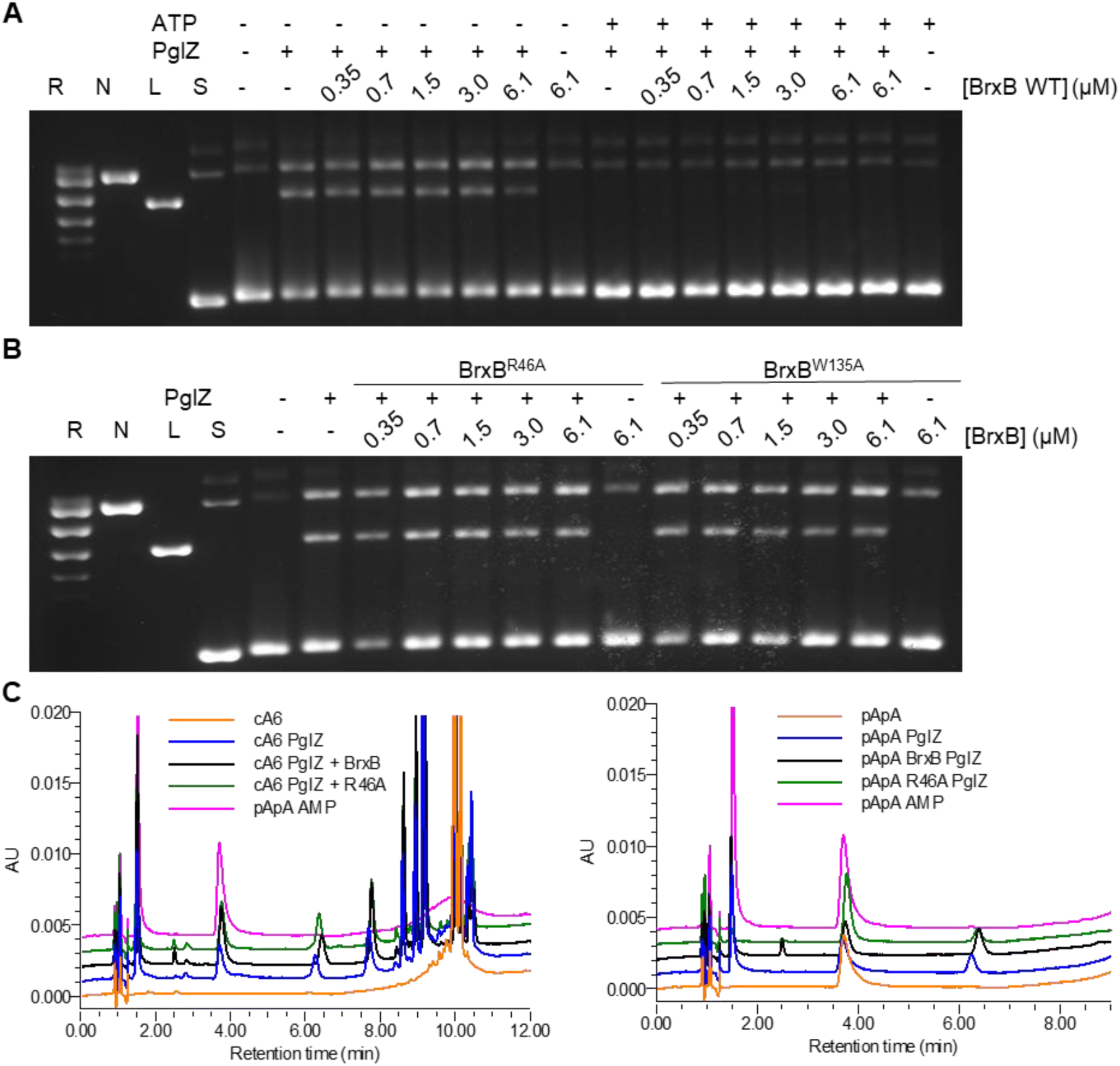
BrxB interacting with PglZ does not appreciably alter PglZ nuclease activity. (**A**) Incubation of PglZ (768 nM) with a titration of BrxB WT in the absence or presence of MnCl_2_ (0.5 mM) and ATP (1 mM). (**B**) Incubation of PglZ WT (768 nM) with a titration of BrxB mutants R46A and W135A in the absence or presence of MnCl_2_ (0.5 mM). Control lanes represent supercoiled (S), nicked (N), linear (L), and relaxed (multiple topoisomers; R) plasmid DNA, or contain DNA in the absence of protein. Assays are presented on 1.4% (w/v) agarose 1x TAE gels post-stained with ethidium bromide. Assays shown are representative of triplicate experiments. (**C**) PglZ (2 μM) incubated in the presence of BrxB (10 μM) does not prevent cleavage of cA6 or pApA (10 μM). Control reactions are comprised of the nucleotide in the absence of protein. Standard mixes are comprised of pApA and AMP. Presented traces are representative of triplicate data.

### PglZ nuclease activity contributes to BREX phage defence but not BREX-dependent methylation

With the *E. fergusonii* PglZ and BrxB mutations now characterised biochemically we examined their impact on the two measurable BREX phenotypes, phage defence and BREX-dependent methylation. Mutations were constructed in the context of pBrxXL, which encodes the full *E. fergusonii* BREX locus under native promoters (24). The suite of mutants were tested for defence against phage Pau from the Durham collection (23), measured by Efficiency of Plating (EOP), using an appropriate vector control. The positive and negative controls pBrxXL and pBrxXL-Δ*pglX* provided strong and no phage defence, respectively (**Fig. 7A**). BrxB mutant constructs S133A and W135A, and the PglZ H741A construct all showed a small reduction in phage defence activity, around 10-fold (**Fig. 7A**). PglZ T538A and double mutant T538A/H741A constructs showed a large reduction in defence of around 3 logs, but remained impressively active (**Fig. 7A**). These data indicate that mutations preventing PglZ:BrxB interactions or ablating PglZ nuclease activity have an impact but can be compensated for *in vivo*.

**Figure 7.**
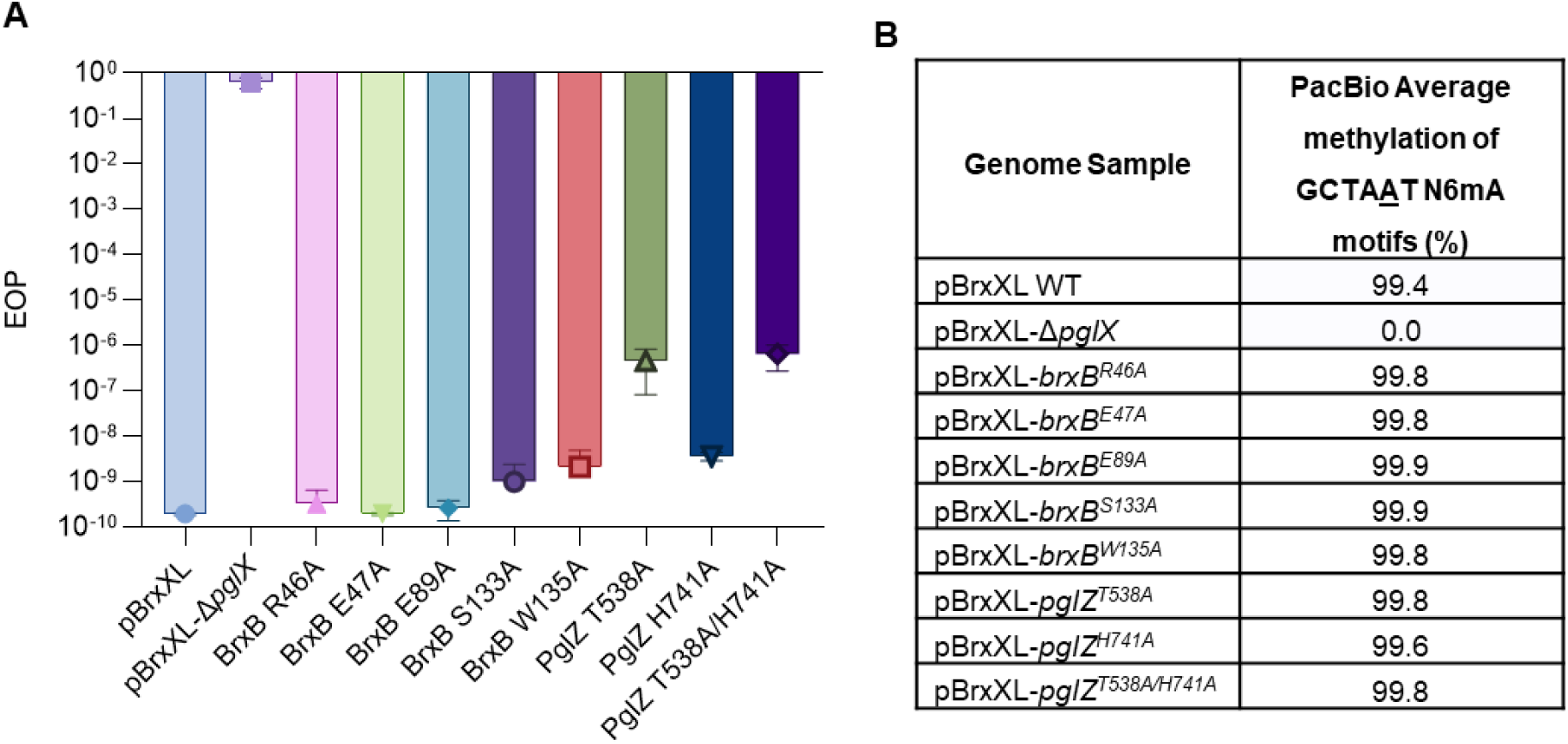
PglZ:BrxB mutants have small impact on BREX phage defence and no impact on BREX methylation. (**A**) EOP results of phage Pau tested against *E. fergusonii* BREX constructs. Error bars represent standard deviation from the mean of triplicate data. (**B**) PacBio sequencing results showing the percentage of BREX motif methylation in each strain.

Genomic DNA was extracted from each strain and PacBio sequencing analysis was performed to investigate BREX-dependent methylation, examining N6mA methylation on the fifth adenine of the GCTAAT motif (**Fig. 7B**). Positive control pBrxXL WT had 99.4% methylation of BREX motifs, and negative control pBrxXL-Δ*pglX* had no detectable methylation, as observed previously (24). In all cases, the current mutations in pBrxXL had no impact on BREX-dependent methylation (**Fig. 7B**), indicating PglZ nuclease activity works independently from BREX methylation.

## Discussion

Type I BREX systems perform phage defence and BREX-dependent methylation, but the mechanisms for each activity have proven difficult to uncover. Having previously performed individual studies of BREX components and regulators (17–22), in this study we examined what higher order BREX complexes form in cells and focussed on characterisation of PglZ, BrxB and the resulting stable PglZ:BrxB sub-complex.

Using *Salmonella* BrxB as bait we observed pull-down of BrxB with BrxC, PglX, PglZ and also BrxL (**Fig. 1**). Although BrxL is the effector protein needed for phage defence with the *E. coli* and *Acinetobacter* BREX systems (20, 39), we previously noted that *Salmonella* BREX can provide phage defence without BrxL (23), and there is potential for BrxL having a regulatory role in this system (40). It was therefore curious that BrxL readily associates with other *Salmonella* BREX components. However, low levels of BrxL can also be seen in similar pull-down experiments performed using *E. coli* BREX (18). Using BrxL as bait does also pull-down other BREX components, though the data do show that the more robust complex appears to be formed of BrxB, BrxC, PglX and PglZ (18). We noted strong association between PglZ and BrxB, also seen with *E. coli* BREX (18), and so chose to examine this sub-complex as a step towards understanding the structure and function of higher order BREX complexes.

Our resulting hybrid model of *Salmonella* PglZ:BrxB (**Fig. 2**) uses 4.45 Å resolution density obtained by cryoEM to fit AlphaFold (33) predicted domains of each protein. Notably, our model demonstrates points of structural flexibility that allow movement of the PglZ:BrxB interaction domain and the PglZ C-terminal β-barrel domain (**Supplementary Fig. S3**). This flexibility is the likely cause of the limited resolution observed for the corresponding cryoEM analysis.

The first evidence of biochemical activity for PglZ domains, originally considered an alkaline phosphatase (15), came from demonstration that the PglZ domain of PorX, a two-component signalling system response regulator, could act as a phosphodiesterase and linearise cyclic nucleotides (16). PorX activity is zinc-dependent. Having examined the structure of *Salmonella* PglZ:BrxB, we switched to *E. fergusonii* PglZ and BrxB for biochemical characterisation as the proteins behaved more reproducibly under assay conditions. After substantial efforts to identify a preferred metal and concentration, it could be demonstrated that *E. fergusonii* PglZ has similar activity to PorX, cleaving a wide range of cyclic nucleotides, but not cyclic mononucleotides or dinucleotide polyphosphates (**Fig. 3 and Supplementary Fig. S4-8**).

We hypothesised that our observed *E. fergusonii* PglZ activity could impact dsDNA. When tested, *E. fergusonii* PglZ nicked and linearised dsDNA (**Fig. 4**). This activity was applicable to ssDNA, but not dsRNA, and was independent of BREX motifs, appeared sequence-independent, was not impacted by BREX methylation and was not impacted by larger DNA modifications such as glucose modifications to hydroxymethylated cytosines in phage T4 genomic DNA (**Fig. 4 and Supplementary Figs. S9 and S10**). Initial investigations of BREX activity indicated little digestion of invading phage DNA, merely an inhibition of phage genome replication, making BREX a classic “restriction” system (15). Nicking is a hallmark of multiple other phage defence systems, including Shedu, Lamassu, Dnd and Gabija (41–45). In the latter case, Gabija activity is also regulated by the detection and degration of nucleotides (45). As we observed inhibition of PglZ nicking activity in the presence of ATP (**Fig. 4**), we cannot rule out analogous regulatory activity by PglZ in the context of a full BREX mechanism. It is unclear whether ATP might be competing for the PglZ domain catalytic site, or have another inhibitory binding site that alters activity. We also cannot dismiss a potential role for PglZ-dependent nicking in protecting from invading DNA, perhaps as a precursor licensing step to allow further inhibition of replication.

The role of BrxB has thus far remained hypothetical, yet the predicted fold mimics AAA+ nucleotide binding domains. We were able to demonstrate binding of *E. fergusonii* BrxB to PglZ, and pinpoint residues required for stable complex formation (**Fig. 5 and Supplementary Fig. S12 and S13**). When we then tested whether BrxB might bind and therefore alter the observed inhibition of PglZ by ATP, nothing changed (**Fig. 6**). Alternatively, and due to the observed data in *Salmonella* (**Fig. 1**) and *E. coli* (18), we postulate that the role of BrxB might be as a scaffold protein, participating within and allowing connections between multiple BREX components within higher order complexes.

When assaying the two phenotypes of phage defence and methylation we saw that ablation of PglZ:BrxB interactions made only a small impact on phage defence (**Fig. 7A**), perhaps because within a higher order complex other interactions occur to support function. In contrast, removal of PglZ nuclease activity by mutation had a stronger impact though still did not remove all phage defence activity, indicating that the overall BREX mechanism can compensate for loss or reduction in PglZ function (**Fig. 7A**). Whilst deletion of *pglZ* prevents BREX phage defence and methylation (17, 20, 39), our mutations of either PglZ nuclease activity or BrxB binding had no impact on methylation (**Fig. 7B**), indicating PglZ likely plays a role in formation of the BREX methylation complex, but not in that specific activity.

A working model for BREX activity was recently posited, wherein a BREX-BCXZ complex would form and move along DNA to allow methylation (18). Our data support formation of this complex (**Fig. 1**), and contrary to data indicating PglX alone is sufficient for methylation (46), we could not see methylation using *Salmonella* or the same *E. coli* homologue (17, 18)and expression of PglX alone in cells also does not result in methylation (39). A BREX-BCXZ complex would have to be able to distinguish between DNA templates containing BREX methylation on one strand following host genome replication, and target invading DNA containing no BREX methylation. When invading DNA is recognised, we envision a role for the PglZ nuclease in which nicking of target DNA licenses the BREX-BCXZ complex to switch from methylation surveillance to restriction. This could include recruitment of BrxL and, for instance, movement of the BREX complex to stall replication forks. Nevertheless, there remains many questions as to the specifics of BREX activity and our data indicate clear next steps in the characterisation of higher order BREX complexes.

## Acknowledgements

This research was supported by the Electron Microscopy Shared Resource, <u>RRID:SCR_022611</u>, of the Fred Hutch/University of Washington/Seattle Children’s Cancer Consortium (P30 CA015704). CryoEM molecular graphics and analyses were performed with UCSF ChimeraX, developed by the Resource for Biocomputing, Visualization, and Informatics at the University of California, San Francisco, with support from National Institutes of Health R01-GM129325 and the Office of Cyber Infrastructure and Computational Biology, National Institute of Allergy and Infectious Diseases.

## Author contributions

J.J.R. and L.A.D. contributed equally. J.J.R. expressed all *E. fergusonii* proteins and performed biochemical analyses. L.A.D. performed cryoEM practical aspects, data collection and data processing. M.P. produced *Salmonella* proteins for biochemistry and performed co-expression analyses and phage assays. A.K. performed *in vivo* pull-down analyses and mass photometry. A.N. performed PacBio sequencing and analysis. A.J.K., S.M. and J.P-A. produced and analysed *Acinetobacter* proteins. T.R.B. produced *Salmonella* protein for cryoEM. D.L.S., B.L.S., B.K.K., and T.R.B. supervised the project and obtained funding. All authors contributed to data analysis and writing the manuscript.

## Supplementary data

Supplementary data have been provided and comprise 13 Figures and one Table.

## Conflict of interest

T.R.B. is an employee of, and B.L.S. is a paid consultant for, New England Biolabs, which provided funding support for this study and which develops a wide variety of phage restriction systems for commercial sale.

## Funding

This work was supported by a Biotechnology and Biological Sciences Research Council Newcastle-Liverpool-Durham Doctoral Training Partnership studentship [grant number BB/T008695/1] to J.J.R., a Biotechnology and Biological Sciences Research Council responsive mode grant [grant number BB/Y003659/1] to M.P., a Lister Institute Prize Fellowship to A.K. and T.R.B., New England Biolabs (NEB), the Fred Hutchinson Cancer Center (FHCC) and the NIH for both BLS (R01 GM105691) and BKK (R15 GM140375).

For the purpose of open access, the authors have applied a CC BY public copyright licence to any Author Accepted Manuscript version arising from this submission.

## Data Availability

The cryoEM model and corresponding maps for PglZ complexed with BrxB have been deposited in the RCSB PDB database (ID code 9NV3) and in the EMDB (ID code EMD-49827). All other data needed to evaluate the conclusions in the paper are present in the paper and/or Supplementary Data.

