## Supplementary Information for "PglZ from Type I BREX phage defence systems is a metal-dependent nuclease that forms a sub-complex with BrxB"

[*]These authors contributed equally.

### Supplementary Figure S1

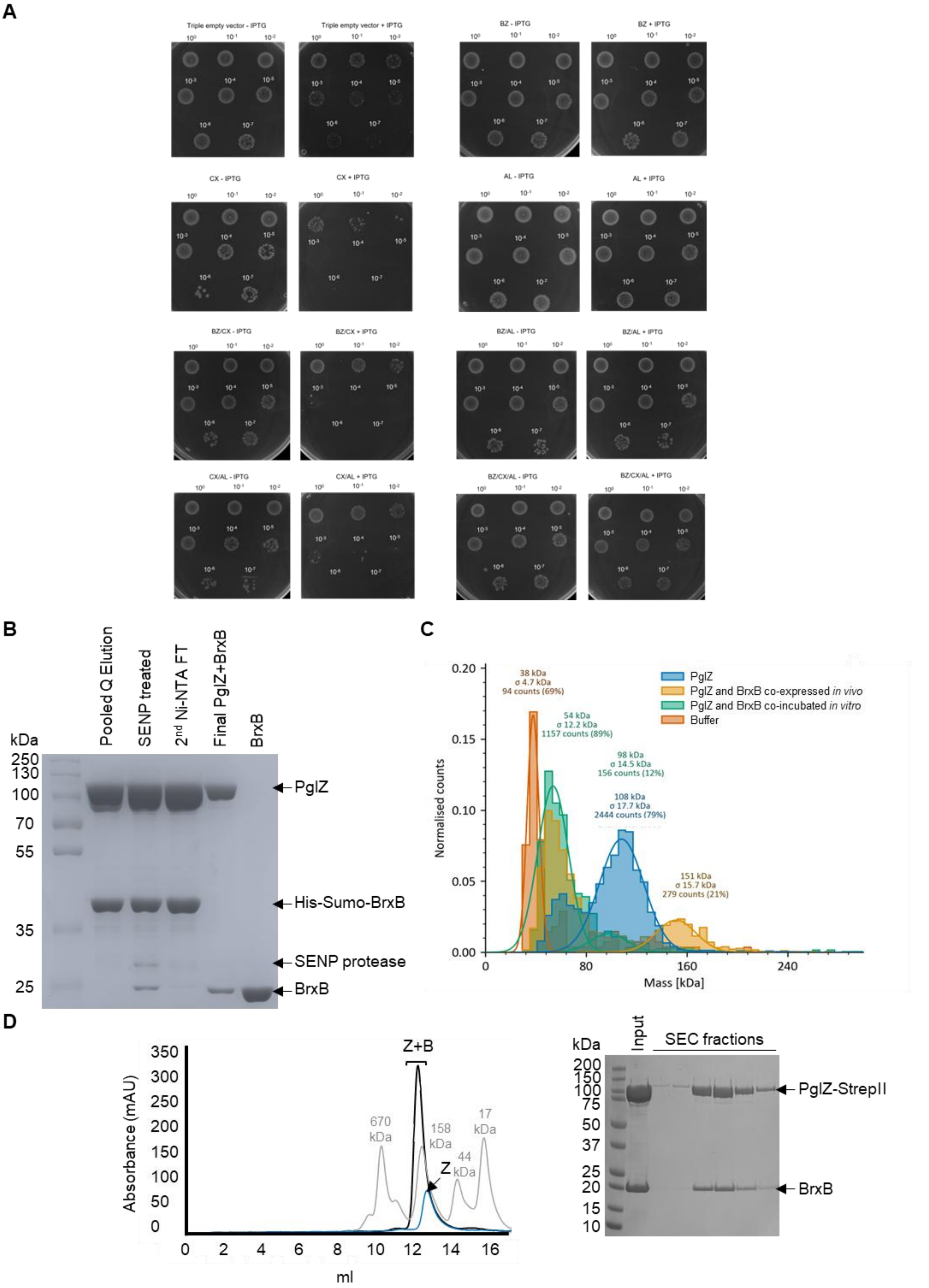


**Supplementary Figure S1.** BREX protein toxicity tests and purification of PglZ:BrxB complexes. (**A**) Spot tests for toxicity during expression of *Salmonella* BREX proteins. Plates are representative of triplicate data. (**B**) Selected samples during production of *Salmonella* PglZ:BrxB complexes for structural study. (**C**) Mass photometry analysis of *Salmonella* PglZ:BrxB samples. (**D**) Size exclusion chromatography traces for *Acinetobacter* PglZ (blue) and PglZ:BrxB (black), with molecular weight standards shown in grey. Fractions for the PglZ:BrxB purification are shown separated by SDS-PAGE.

**Supplementary Figure S2**

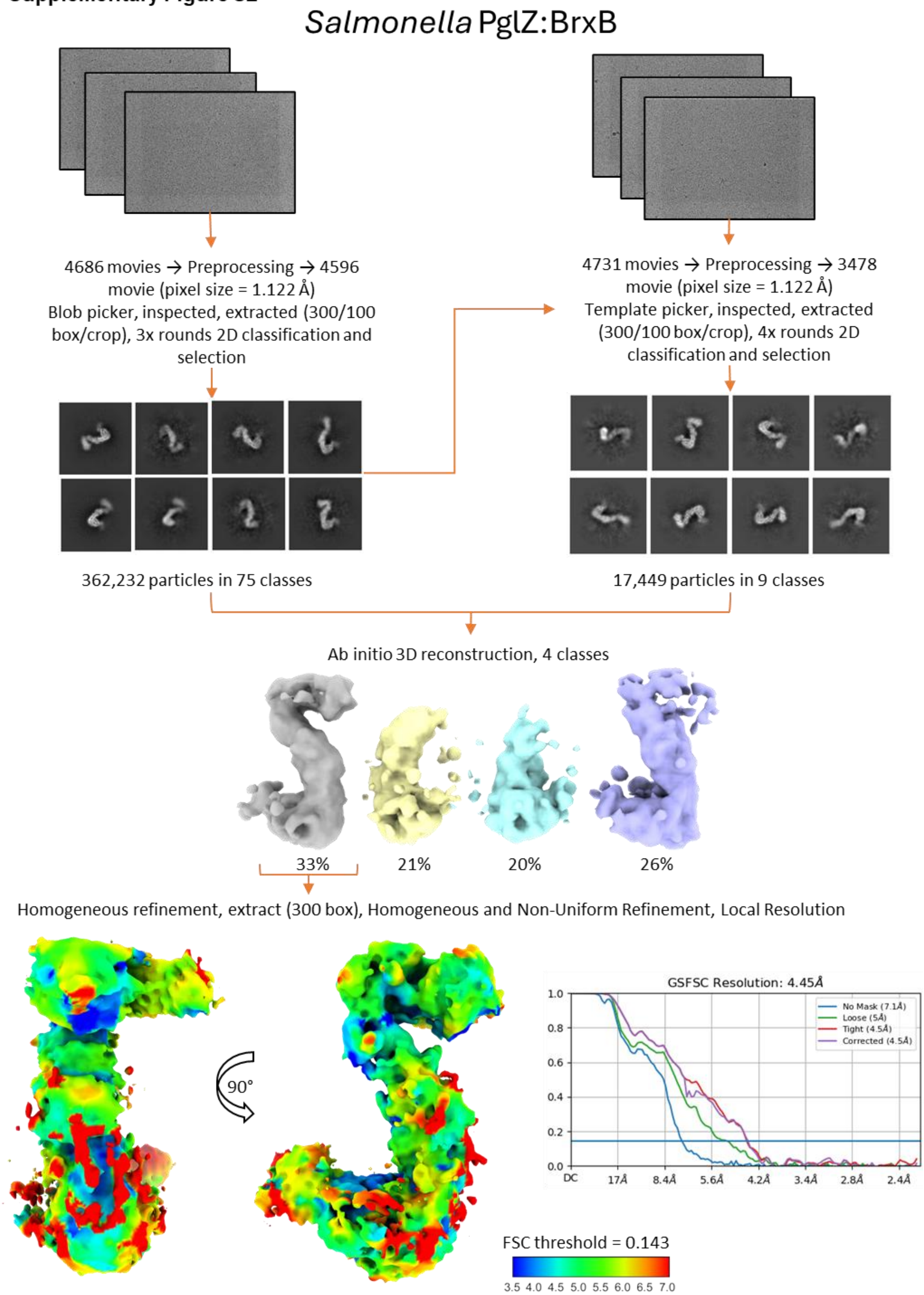


**Supplementary Figure S2.** Workflow for processing of the *Salmonella* PglZ:BrxB sub-complex data.

**Supplementary Figure S3**

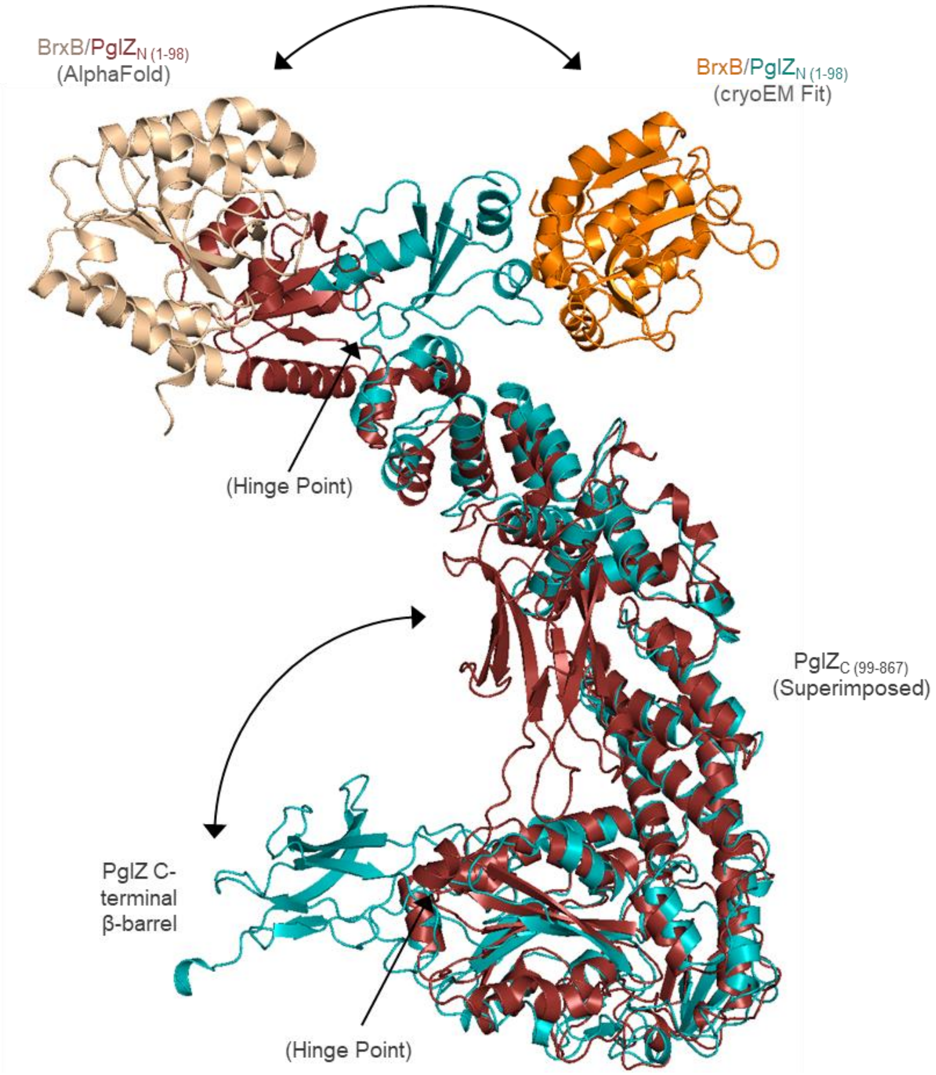


**Supplementary Figure S3.** *Salmonella* PglZ has a flexible N-terminal domain that allows movement of BrxB, and a second hinge point that alters position of the C-terminal β-barrel domain.

**Supplementary Figure S4**

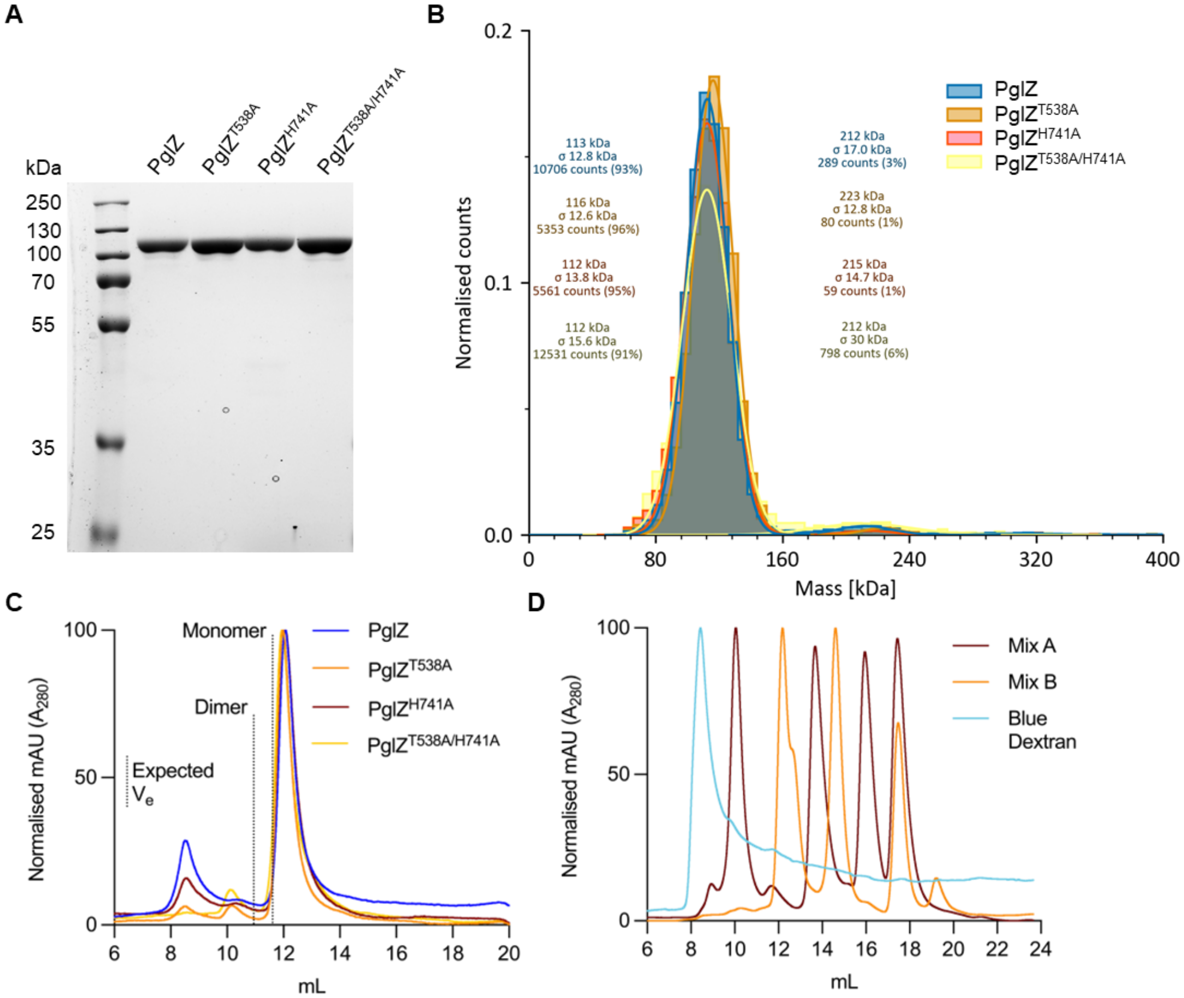


**E**

| | Calculated Rst (Å) | Observed Rst (Å) | Observed Rst (Å)/ Calculated Rst (Å) |
|---|---|---|---|
| Monomer | 50.11 | 46.70 | 0.93 |
| Dimer | 56.00 | 62.46 | 1.12 |

**Supplementary Figure S4.** *E. fergusonii* PglZ WT and mutant production. (**A**) SDS-PAGE of PglZ WT and mutants loaded at 4 µg on 15% (v/v) polyacrylamide gel. Samples were resolved for 1 h 15 min in tris-glycine running buffer and stained with Quick Coomassie. (**B**) Mass photometry of PglZ WT and mutants. Counts were acquired for 60 s with 5 nM protein samples in PBS. (**C**) Analytical SEC (S200i) of PglZ WT and mutants. Protein samples were loaded at 10 µM in a 100 µl final volume. The expected elution volumes ($V_e$) of monomeric and dimeric PglZ are represented by black dotted lines. (**D**) Calibration of analytical SEC S200i column using protein standards. Mix A: RNase A, Ferritin, Conalbumin, and Carbonic anhydrase. Mix B: RNase A, Aldolase, Ovalbumin, and Aprotinin. (**E**) Comparison of expected elution points based on Stokes radii. Presented data are representative of triplicate data.

**Supplementary Figure S5**

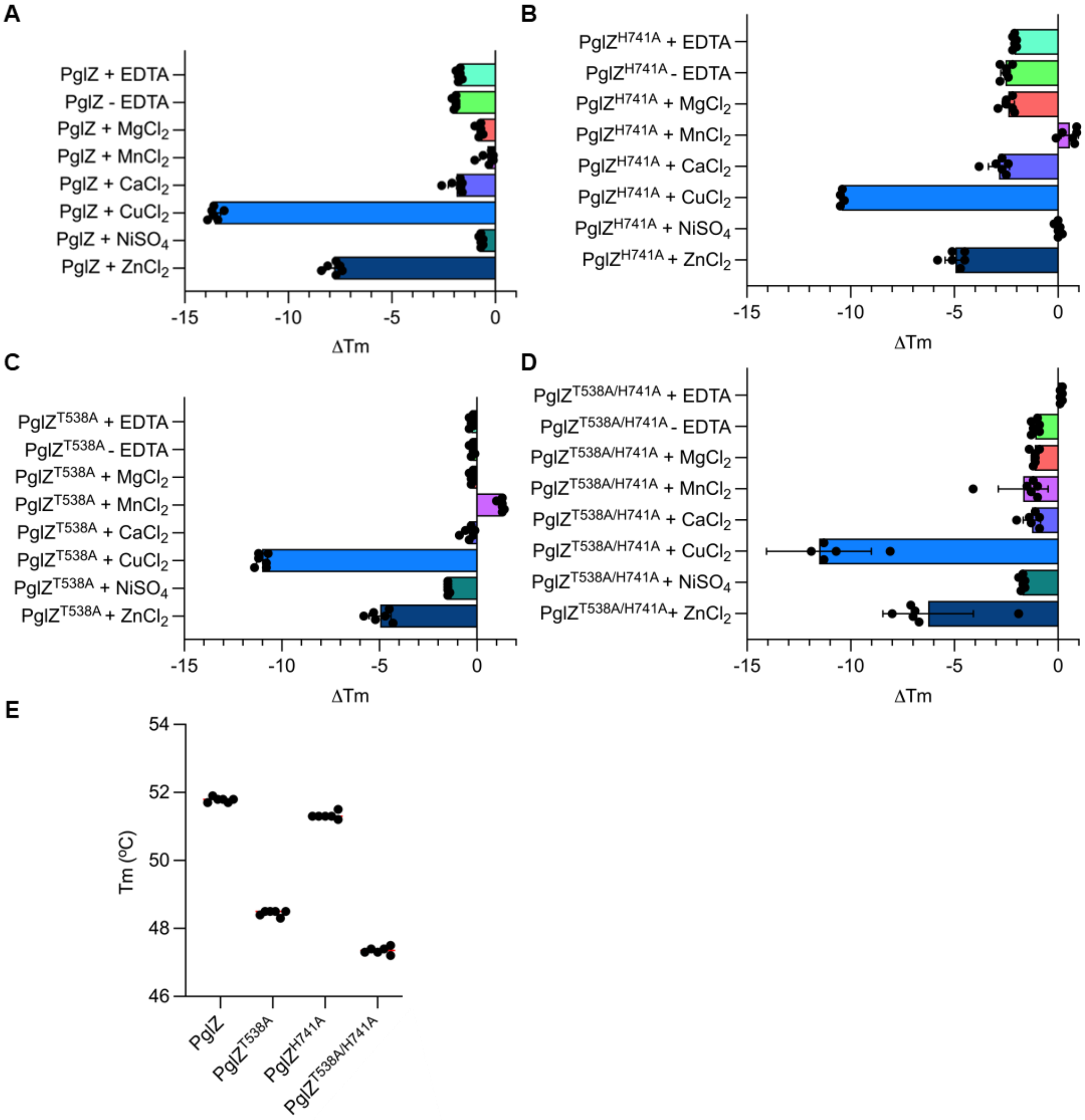


**Supplementary Figure S5.** Thermal shift assay of *E. fergusonii* PglZ WT (**A**) and mutants H741A (**B**), T538A (**C**), and T538A/H741A (**D**), showing metal-dependent thermostability. Mean changes in melting temperature (ΔTm) are plotted by comparison to respective PglZ WT or mutants in the absence of EDTA or metal, which is set at point '0'. (**E**) The melting temperature (Tm) of PglZ WT and mutants. Protein samples were analysed at 5 μM. Plotted data represent the mean ± SD (6 replicates).

### Supplementary Figure S6

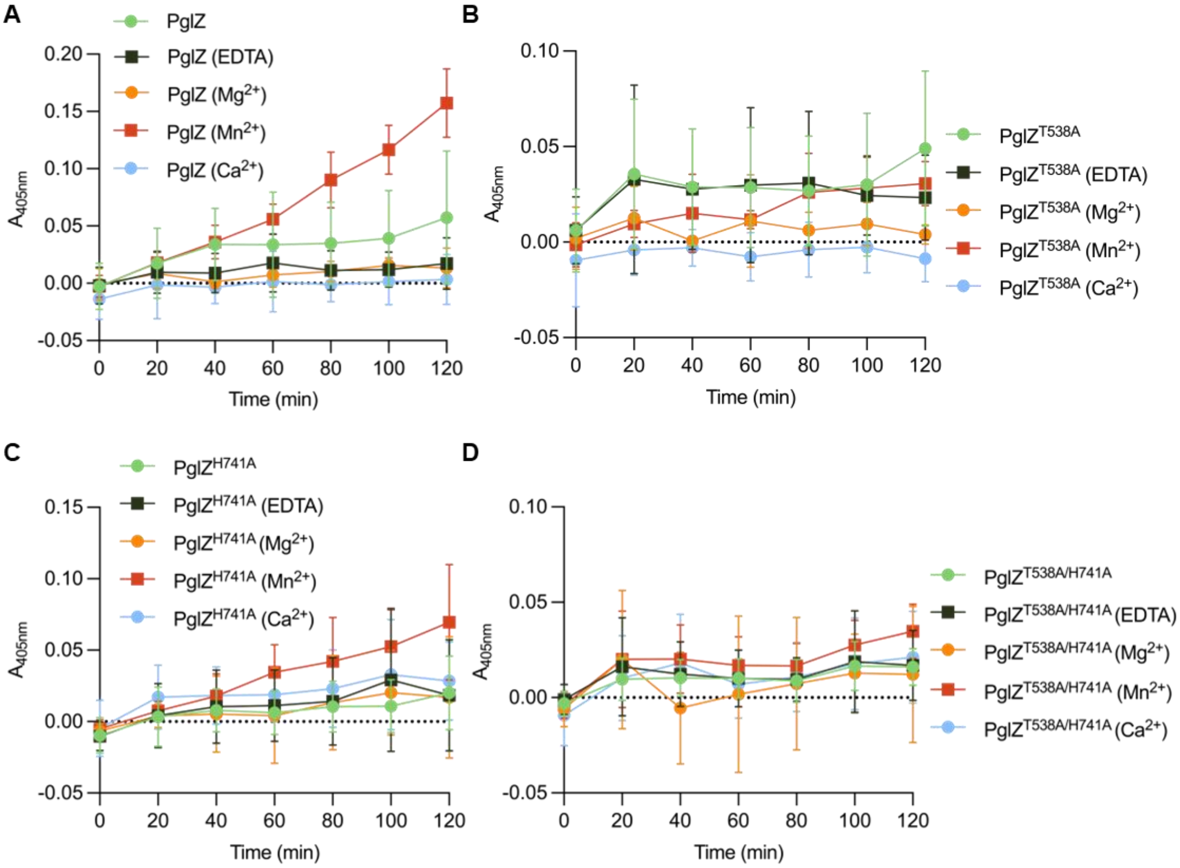


**Supplementary Figure S6.** *E. fergusonii* PglZ WT (**A**) and mutants T538A (**B**), H741A (**C**), and T538A/H741A (**D**) Bis-*p*NPP phosphodiesterase assays, showing metal-dependent phosphodiesterase activity. Absorbance ($A_{405nm}$) represents the amount of reaction product *p*-nitrophenyl phosphate. Protein samples were tested at 2 µM in the presence of 2.5 mM Bis-*p*NPP, with or without EDTA and 0.5 mM metal. Control reactions comprised of metal buffer and Bis-*p*NPP in the absence of PglZ, which were set at absorbance '0' for each timepoint. Plotted data represent mean absorbance readings ± SD (9 replicates).

### Supplementary Figure S7

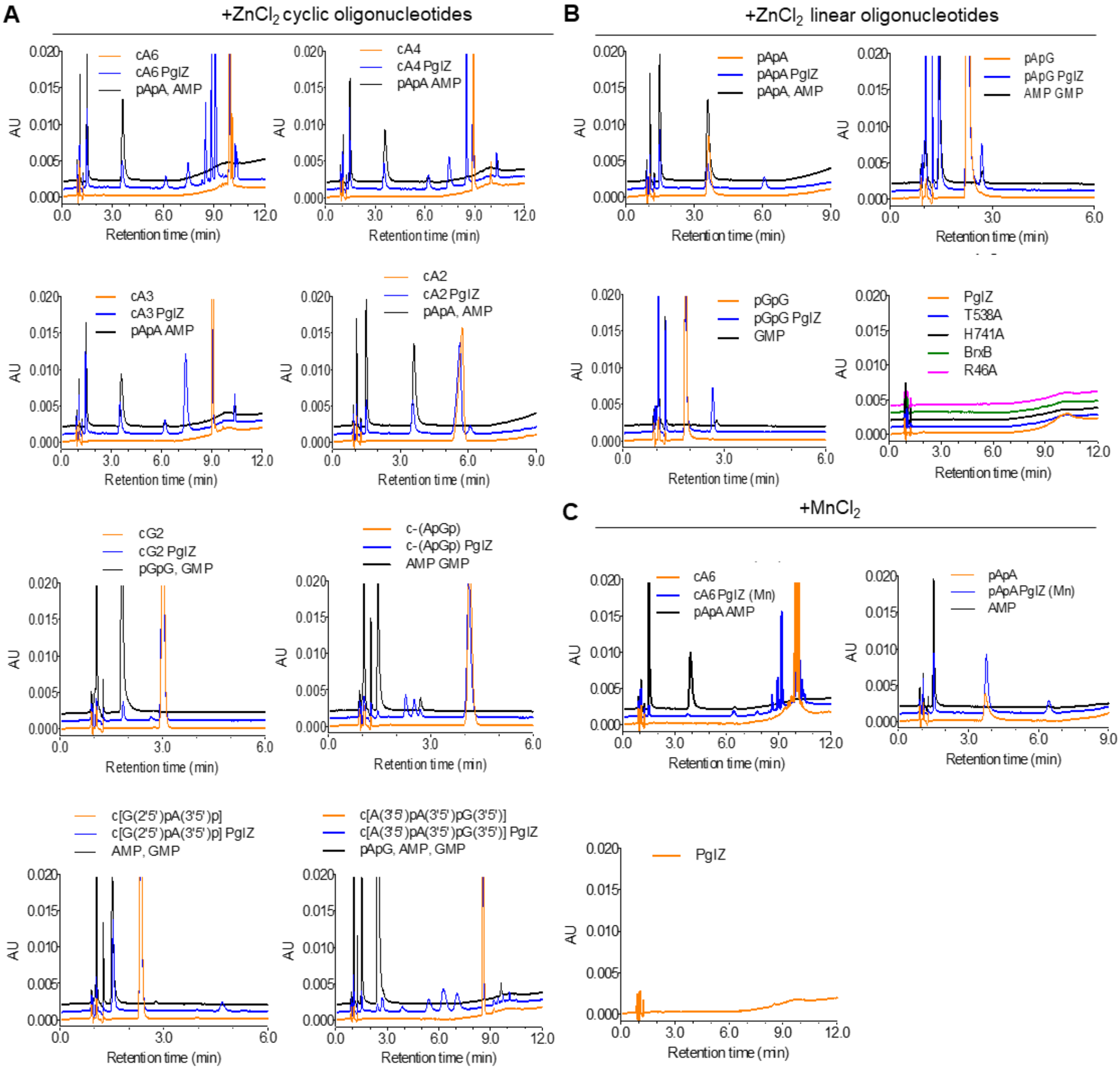


**Supplementary Figure S7.** *E. fergusonii* PglZ (2 µM) can cleave a range of A and G nucleotides (10 µM) in a zinc- or manganese-dependent manner. (**A**) HPLC analysis of PglZ activity on cyclic oligonucleotides, using $ZnCl_2$ (10 µM). (**B**) HPLC analysis of PglZ activity on linear oligonucleotides, using $ZnCl_2$ (10 µM). (**C**) HPLC analysis of PglZ activity on cA6 and pApA, using $MnCl_2$ (10 µM). Control reactions are comprised of the nucleotide in the absence of protein, and the protein in the absence of nucleotide. Standard mixes are comprised of expected degradation products for each reaction. Presented traces are representative of triplicate data.

### Supplementary Figure S8

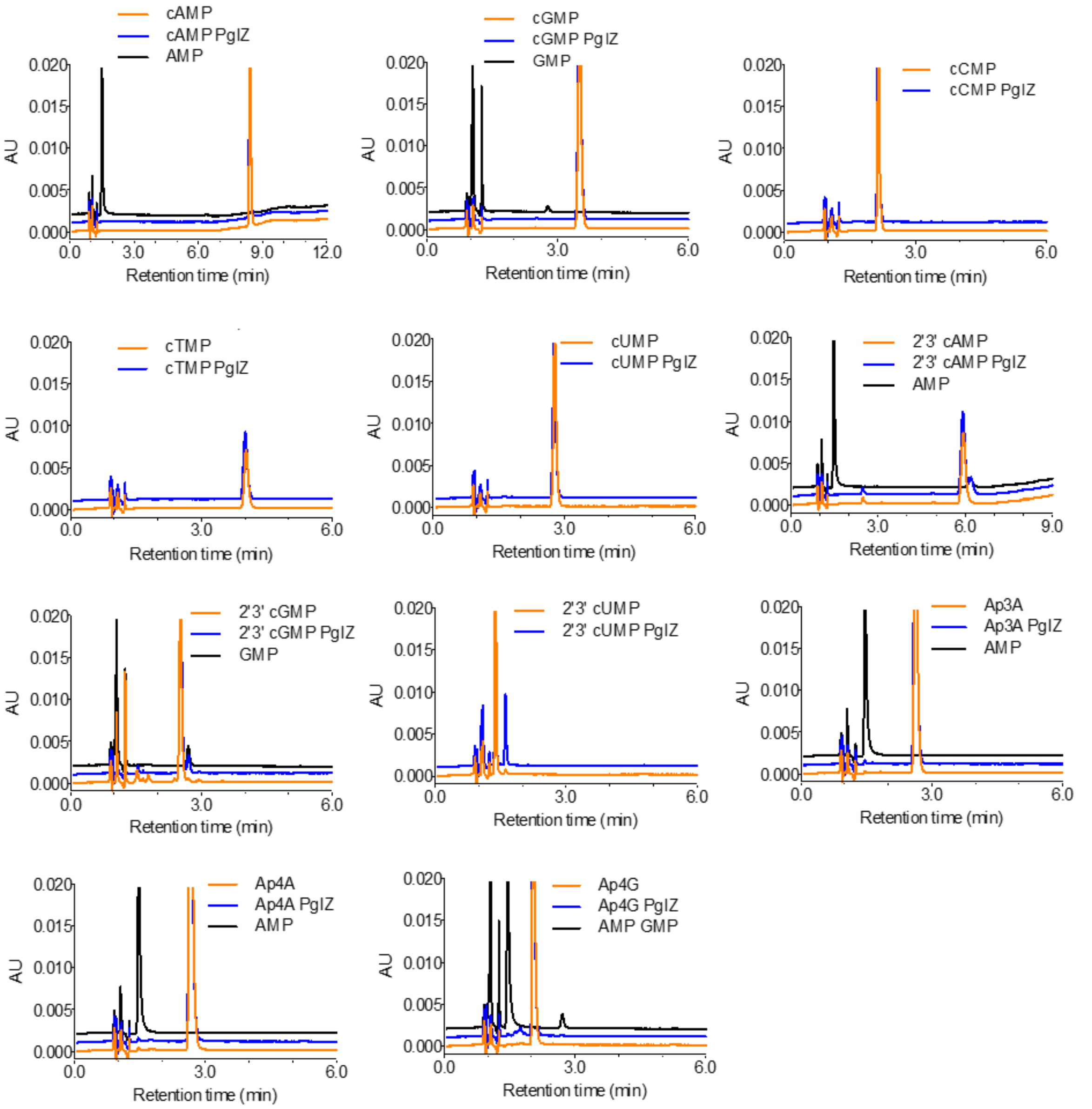


**Supplementary Figure S8.** *E. fergusonii* PglZ (2 µM) has no cleavage activity on 3′-5′ or 2′-3′ cyclic mononucleotides or dinucleotide polyphosphates (10 µM) in the presence of $ZnCl_2$ (10 µM), analysed by HPLC. Control reactions are comprised of the nucleotide in the absence of protein. Standard mixes are comprised of expected degradation products for each reaction, where possible. Presented traces are representative of triplicate data.

### Supplementary Figure S9

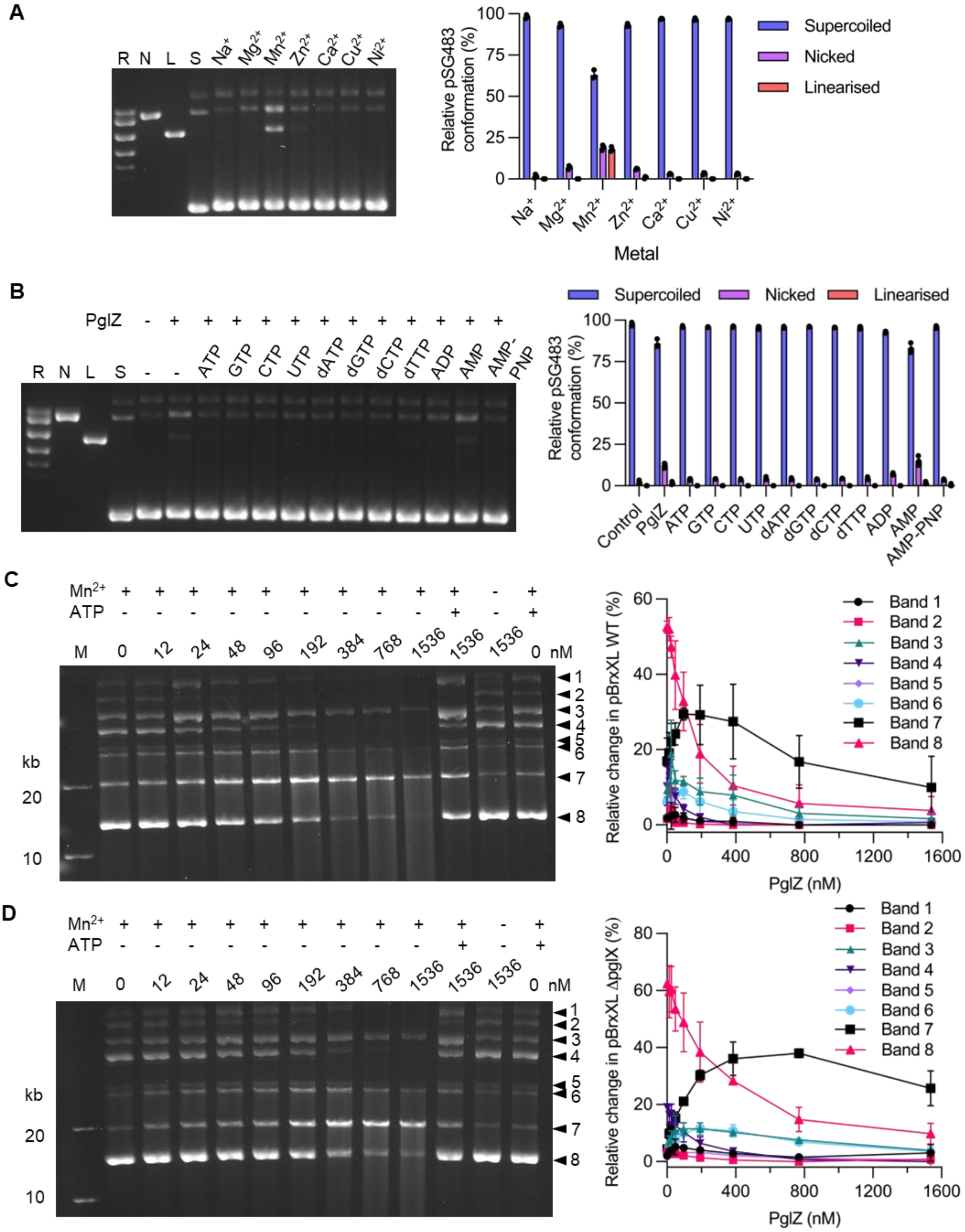


**Supplementary Figure S9.** Impact of metals, nucleotides, and BREX methylation on *E. fergusonii* PglZ nuclease activity. (**A**) PglZ nuclease activity is maximised in vitro by Mn. PglZ (768 nM) was tested in the presence of 0.5 mM of various metals against supercoiled pSG483 DNA (6 nM) (**B**) A wide range of nucleotides (1 mM) can inhibit PglZ (768 nM) nuclease activity on supercoiled pSG483 DNA (6 nM). PglZ cleavage of pBrxXL WT (**C**) and pBrxXL-Δ*pglX* (**D**) plasmids, which are BREX methylated and non-methylated, respectively. PglZ was titrated against pBrxXL WT (200 ng) and pBrxXL-Δ*pglX* (200 ng) in the presence and absence of $MnCl_2$ (0.5 mM) and ATP (1 mM). Control lanes represent supercoiled (S), nicked (N), linear (L), and relaxed (multiple topoisomers; R) plasmid DNA, or contain DNA in the absence of protein. Assays are presented on 1.4% (w/v) or 0.8% (w/v) agarose 1x TAE gels post-stained with ethidium bromide. Assays shown are representative of triplicate experiments. Data points and error bars represent the mean ± SD of triplicate data.

**Supplementary Figure S10**

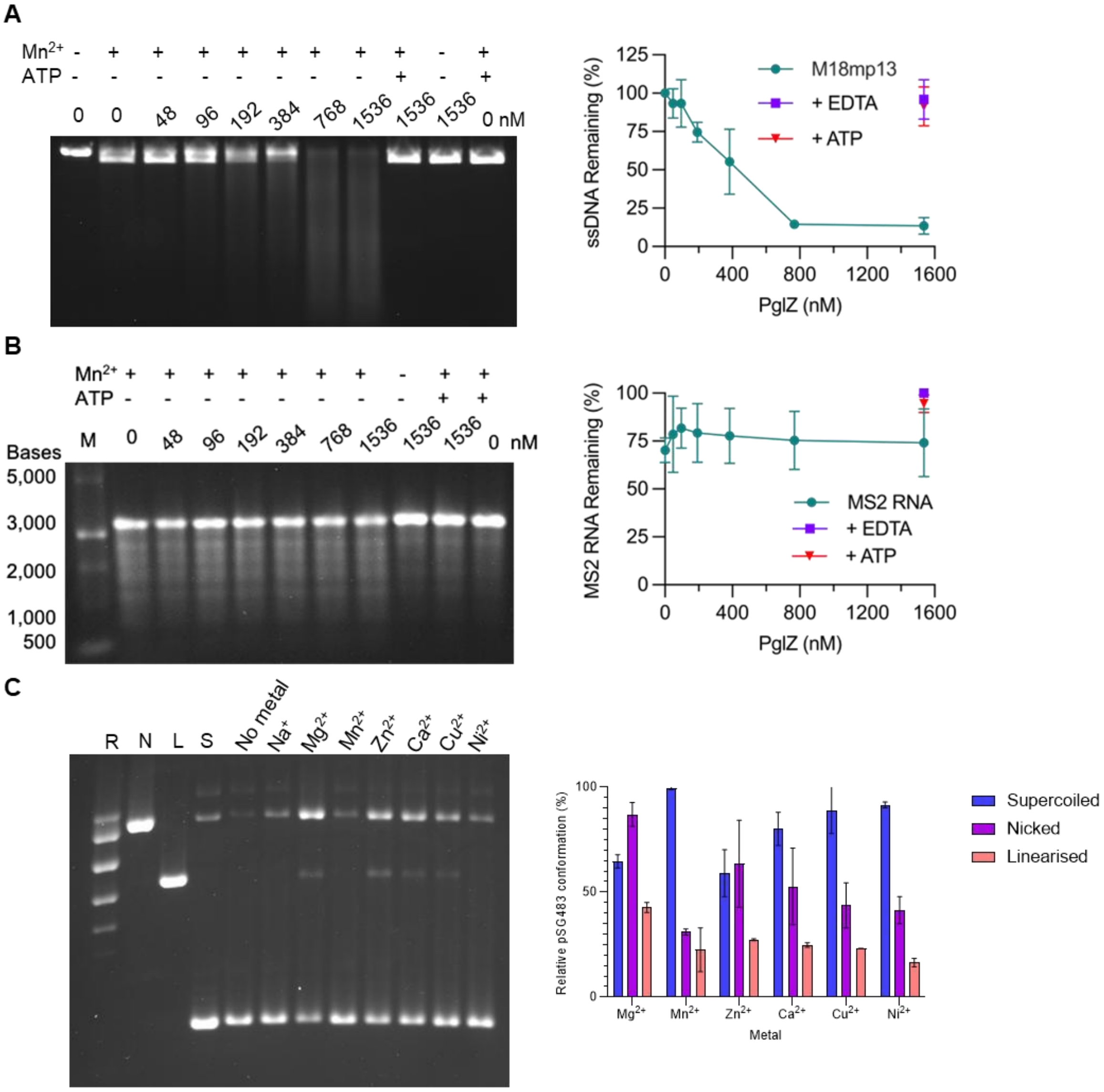


**Supplementary Figure S10.** *E. fergusonii* PglZ can cut a range of nucleic acids, and *Salmonella* PglZ also demonstrates nuclease activity. (**A**) PglZ (768 nM) cleaves M18mp13 ssDNA (6 nM) in the presence of Mn (0.5 mM). (**B**) PglZ (768 nM) is not able to cleave phage MS2 RNA (6 nM). (**C**) *Salmonella* PglZ nicks and linearises supercoiled pSG483 DNA (6 nM) in a metal-dependent manner. Metal was supplied at 0.5 mM. Control lanes represent supercoiled (S), nicked (N), linear (L), and relaxed (multiple topoisomers; R) plasmid DNA, or contain DNA in the absence of protein. Assays are presented on 1.4% (w/v) or 0.8% (w/v) agarose 1x TAE gels post-stained with ethidium bromide. Assays shown are representative of triplicate experiments. Data points and error bars represent the mean ± SD of triplicate data.

### Supplementary Figure S11

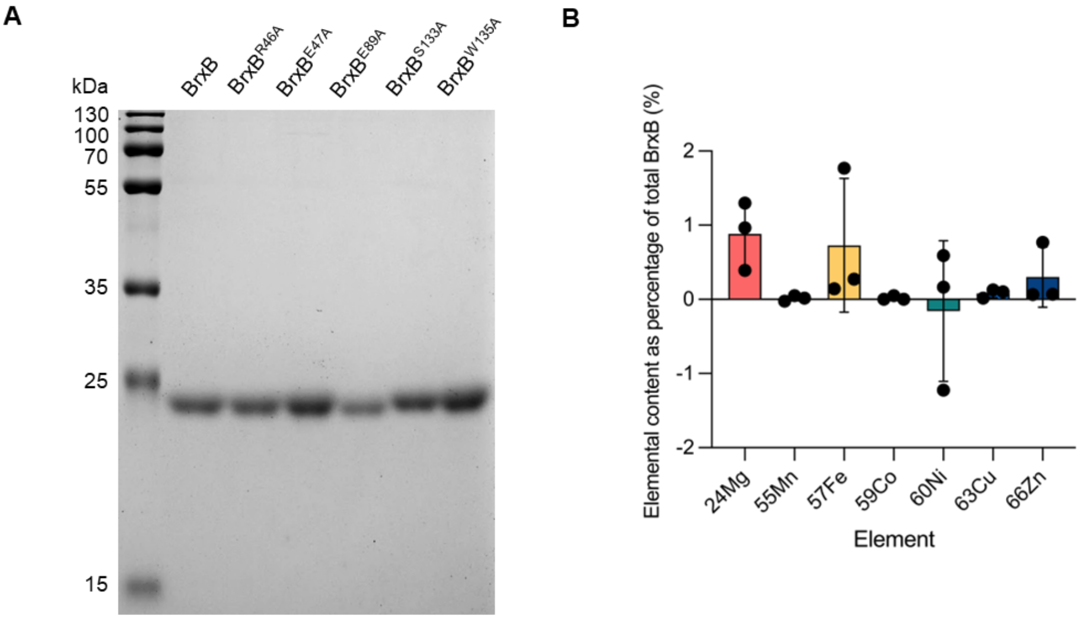


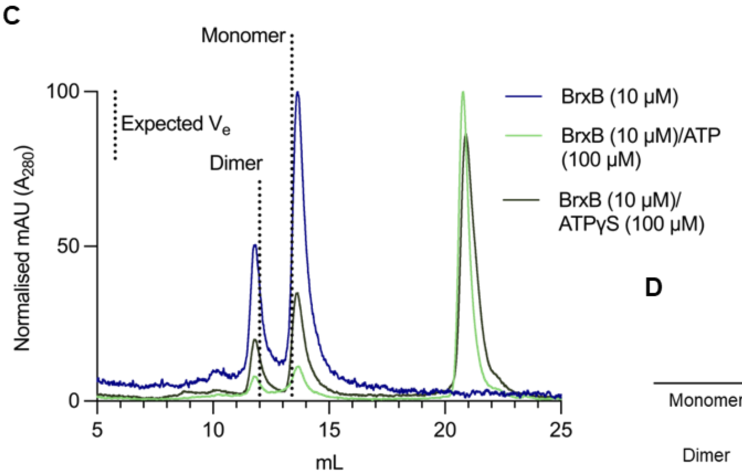


**D**

| | Calculated Rst (Å) | Observed Rst (Å) | Observed Rst (Å)/ Calculated Rst (Å) |
|---|---|---|---|
| Monomer | 23.7 | 23.3 | 0.98 |
| Dimer | 29.1 | 30.2 | 1.04 |

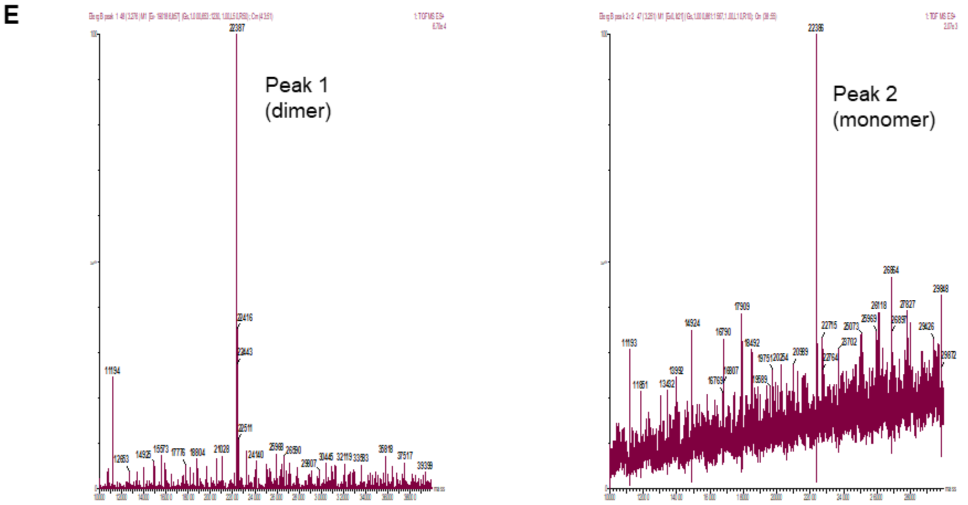


**Supplementary Figure S11.** *E. fergusonii* BrxB does not bind metals and exists as both dimers and monomers. (**A**) SDS-PAGE of purified *E. fergusonii* BrxB WT and mutants loaded at 4 µg on a 12% (v/v) polyacrylamide gel. Samples were resolved for 1 h 15 min in tris-glycine running buffer and stained with Quick Coomassie. (**B**) ICP-MS of BrxB WT and mutants showing divalent cations bound following purification. Plotted data represent mean values ± SD. Metal content is plotted as a percentage of the total protein in the sample. (**C**) Analytical SEC (S75i) of BrxB WT (10 µM) with and without nucleotides and comparison against expected elution ($V_e$) using Stokes radii. (**D**). The expected elution volumes ($V_e$) of monomeric and dimeric BrxB are represented by black dotted lines. (**E**) Electrospray ionisation mass spectrometry of 'monomeric' and 'dimeric' SEC peaks shows BrxB is present in both peaks.

### Supplementary Figure S12

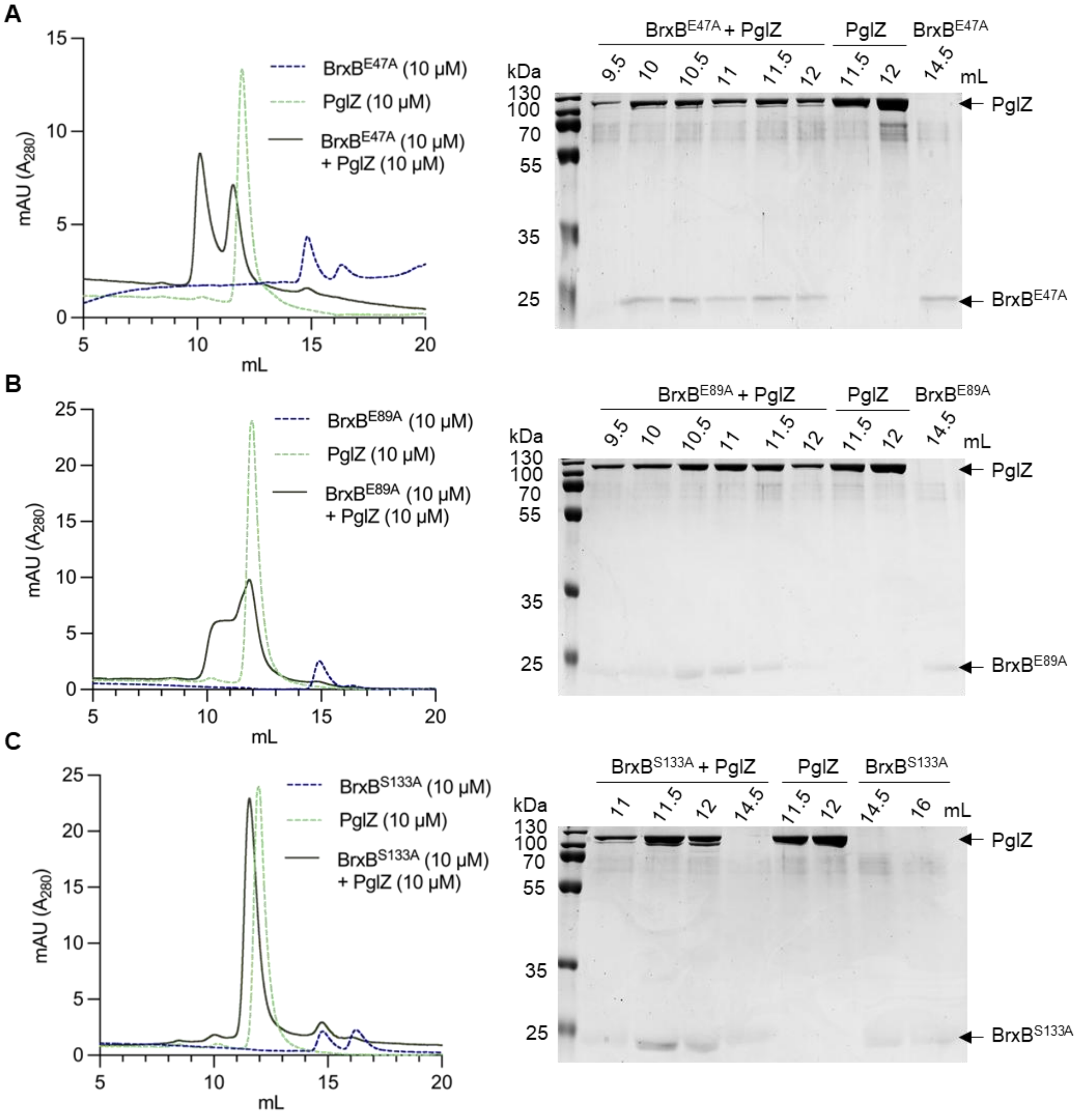


**Supplementary Figure S12.** Analytical SEC (S200i) and SDS-PAGE analysis of *E. fergusonii* PglZ and BrxB mutants. PglZ forms complexes with BrxB mutants E47A (**A**), E89A (**B**), and S133A (**C**). Samples of PglZ (10 µM) with equimolar BrxB WT and mutants were made up to 100 µl and pre-incubated for 15 min prior to loading on the S200i. The elution profile of PglZ incubated with BrxB is shown as a dark green solid line. Control elution profiles of PglZ alone (light green dashed line) and BrxB alone (dark blue dashed line) are also shown. Fractionated peak samples were resolved on 15% (v/v) polyacrylamide gels for 1 h 15 min in tris-glycine running buffer and stained with Quick Coomassie. Protein identities are highlighted with black arrows.

**Supplementary Figure S13**

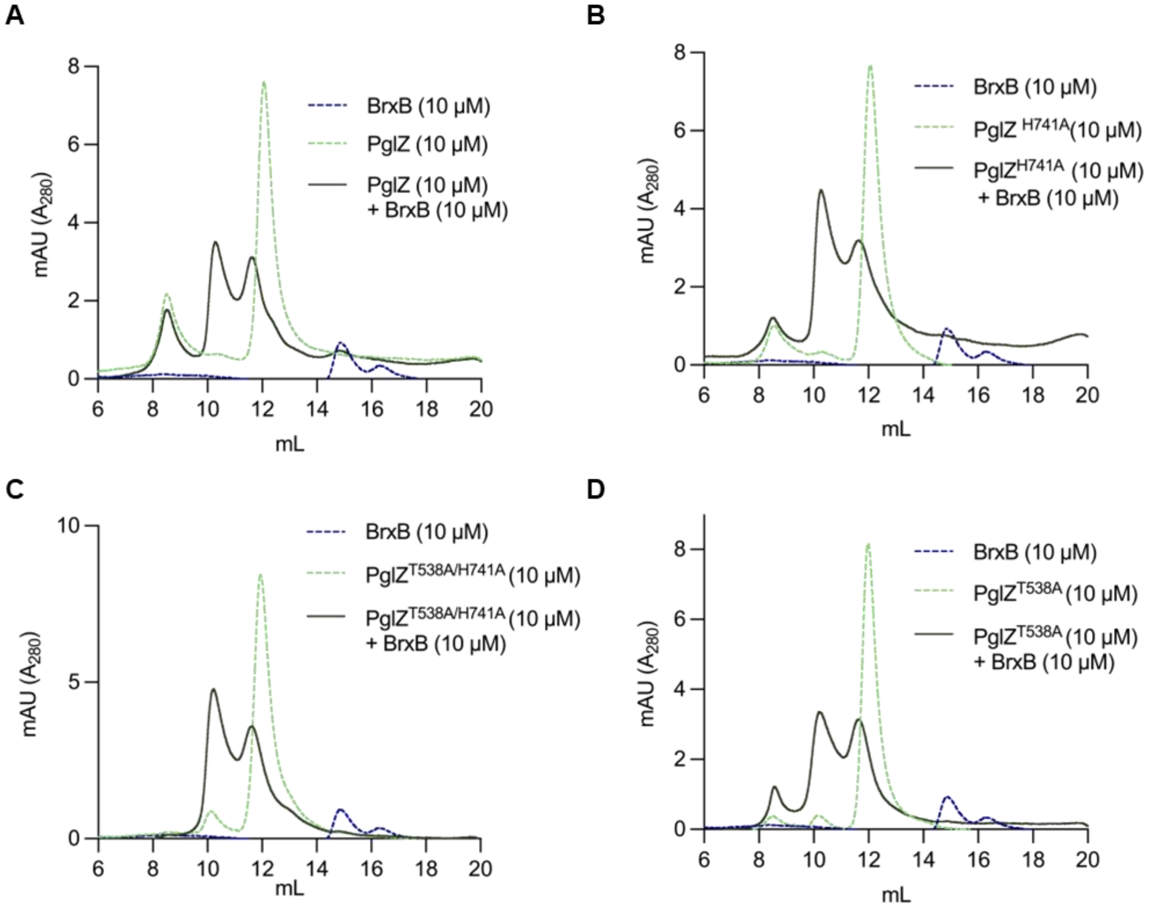


**Supplementary Figure S13.** *E. fergusonii* PglZ WT (**A**) and mutants H741A (**B**), T538A/H741A (**C**), and T538A (**D**) interact similarly with BrxB WT, analysed by analytical SEC (S200i). Samples of BrxB (10 µM) with equimolar PglZ WT and mutants were made up to 100 µl and pre-incubated for 15 min prior to loading on the S200i. The elution profile of PglZ incubated with BrxB is shown as a dark green solid line. Control elution profiles of PglZ alone (light green dashed line) and BrxB alone (dark blue dashed line) are also shown.

**Supplementary Table S1**. Plasmids used in this study.

| PLASMID | NOTES | SYNTHESIS | REFERENCE |
|---|---|---|---|
| p2HR-T | $Ap^R$ | | Addgene #29718 |
| p2HR-T-brxB | *Salmonella* 2HR-T His-strep-BrxB. $Ap^R$ | Genscript | This study |
| pCDF DUET1 | $Sp^R$ | | Novagen |
| pCOLA DUET1 | $Km^R$ | | Novagen |
| pET DUET1 | $Ap^R$ | | Novagen |
| pET15b.PglZ$^{Aci}$-TST | *Acinetobacter* PglZ. C-terminal Twin-Strep tag (thrombin cleavable). $Ap^R$. | Gibson subcloning | This study |
| pET24d.BrxB$^{Aci}$ | *Acinetobacter* BrxB, untagged. $Km^R$ | Gibson subcloning | This study |
| pSALMB | *Salmonella* pSAT-LIC-*brxB*. $Ap^R$ | Genscript | This study |
| pSALMZ | *Salmonella* pSAT-LIC-*pglZ*. $Ap^R$ | Genscript | This study |
| pSALM-BREX | *Salmonella* pCOLA DUET1 *brxA*, *brxB*, *brxC*, *pglX* (MCS1) and *pglZ*, *brxL* (MCS2). $Km^R$ | Genscript | This study |
| pSG483 | $Ap^R$ | | Beck *et al.*, 2024 |
| pSG483-BREX KO | $Ap^R$ | Genscript | This study |
| pTRB444 | *E. fergusonii* pSAT1-LIC-*brxB*. $Ap^R$ | Genscript | This study |
| pTRB473 | *E. fergusonii* pSAT1-LIC-*pglZ*. $Ap^R$ | Genscript | This study |
| pTRB563 | *E. fergusonii* pBrxXL WT. $Cm^R$ | Golden gate assembly | Picton *et al.*, 2021 |
| pTRB564 | *E. fergusonii* pBrxXL-*ΔpglX*. $Cm^R$ | Golden gate assembly | Picton *et al.*, 2021 |
| pTRB710 | *Salmonella* pET DUET1 His-SUMO-*brxB* and *pglZ.* $Ap^R$ | Genscript | This study |
| pTRB724 | *E. fergusonii* pSAT1-LIC-*brxB* E47A. $Ap^R$ | Genscript | This study |
| pTRB725 | *E. fergusonii* pSAT1-LIC-*brxB* S133A. $Ap^R$ | Genscript | This study |
| pTRB726 | *E. fergusonii* pSAT1-LIC-*brxB* R46A. $Ap^R$ | Genscript | This study |
| pTRB727 | *E. fergusonii* pSAT1-LIC-*brxB* W135A. $Ap^R$ | Genscript | This study |
| pTRB728 | *E. fergusonii* pSAT1-LIC-*brxB* E89A. $Ap^R$ | Genscript | This study |
| pTRB729 | *E. fergusonii* pSAT1-LIC-*pglZ* T538A. $Ap^R$ | Genscript | This study |
| pTRB730 | *E. fergusonii* pSAT1-LIC-*pglZ* H741A. $Ap^R$ | Genscript | This study |
| pTRB744 | *E. fergusonii* pBrxXL *brxB* W135A. $Cm^R$ | Genscript | This study |
| pTRB745 | *E. fergusonii* pBrxXL *pglZ* H741A. $Cm^R$ | Genscript | This study |
| pTRB746 | *E. fergusonii* pBrxXL *brxB* E47A. $Cm^R$ | Genscript | This study |
| pTRB747 | *E. fergusonii* pBrxXL *brxB* E89A. $Cm^R$ | Genscript | This study |
| pTRB748 | *E. fergusonii* pBrxXL *brxB* S133A. $Cm^R$ | Genscript | This study |
| pTRB749 | *E. fergusonii* pBrxXL *brxB* R46A. $Cm^R$ | Genscript | This study |
| pTRB750 | *E. fergusonii* pBrxXL *pglZ* T538A. $Cm^R$ | Genscript | This study |
| pTRB758 | *Salmonella* pCOLA DUET1 *brxA* and *brxL.* $Km^R$ | Genscript | This study |

| | | | |
|---|---|---|---|
| pTRB759 | *Salmonella* pCDF DUET1 *brxC* and *pglX*. $Sp^R$ | Genscript | This study |
| pTRB763 | *E. fergusonii* pSAT1-LIC-*pglZ* T538A/H741A. $Ap^R$ | Genscript | This study |
| pTRB766 | *E. fergusonii* pBrxXL *pglZ* T538A/H741A. $Cm^R$ | Genscript | This study |
